# Sex Differences in Neural Networks Recruited by Frontloaded Binge Alcohol Drinking

**DOI:** 10.1101/2024.02.08.579387

**Authors:** Cherish E. Ardinger, Yueyi Chen, Adam Kimbrough, Nicholas J. Grahame, Christopher C. Lapish

**Author notes:** This work was supported in part by grant #s: AA007462, AA023786 (CCL), AA029985 (AK), AA027301 (AK), and the Indiana Alcohol Research Center P60-AA007611. Data Availability Statement: Correlation matrix data are openly available via FigShare: https://figshare.com/articles/dataset/Correlation_Matrix_Data_Sex_Differences_in_Neural_Networks_Recruited_by_Front-loaded_Binge_Alcohol_Drinking_/24867054. Additional data can be made available by request.

## Abstract

Frontloading is an alcohol drinking pattern where intake is skewed toward the onset of access. The goal of the current study was to identify brain regions involved in frontloading. Whole brain imaging was performed in 63 C57Bl/6J (32 female and 31 male) mice that underwent 8 days of binge drinking using the drinking-in-the-dark (DID) model. On days 1-7, three hours into the dark cycle, mice received 20% (v/v) alcohol or water for two hours. Intake was measured in 1-minute bins using volumetric sippers, which facilitated analyses of drinking patterns. On day 8 mice were perfused 80 minutes into the DID session and brains were extracted. Brains were then processed to stain for Fos protein using iDISCO+. Following light sheet imaging, ClearMap2.1 was used to register brains to the Allen Brain Atlas and detect Fos+ cells. For brain network analyses, day 8 drinking patterns were used to characterize mice as frontloaders or non-frontloaders using a recently developed change-point analysis. Based on this analysis the groups were female frontloaders (n = 20), female non-frontloaders (n = 2), male frontloaders (n = 13) and male non-frontloaders (n = 8). There were no differences in total alcohol intake in animals that frontloaded versus those that did not. Only two female mice were characterized as non-frontloaders, thus preventing brain network analysis of this group. Functional correlation matrices were calculated for each group from log_10_ Fos values. Euclidean distances were calculated from these R values and hierarchical clustering was used to determine modules (highly connected groups of brain regions). In males, alcohol access decreased modularity (3 modules in both frontloaders and non-frontloaders) as compared to water drinkers (7 modules). In females, an opposite effect was observed. Alcohol access (9 modules for frontloaders) increased modularity as compared to water drinkers (5 modules). These results suggest sex differences in how alcohol consumption reorganizes the functional architecture of neural networks. Next, key brain regions in each network were identified. Connector hubs, which primarily facilitate communication between modules, and provincial hubs, which facilitate communication within modules, were of specific interest for their important and differing roles. In males, 4 connector hubs and 17 provincial hubs were uniquely identified in frontloaders (i.e., were brain regions that did not have this status in male non-frontloaders or water drinkers). These represented a group of hindbrain regions (e.g., locus coeruleus and the pontine gray) functionally connected to striatal/cortical regions (e.g., cortical amygdalar area) by the paraventricular nucleus of the thalamus. In females, 16 connector and 17 provincial hubs were uniquely identified which were distributed across 8 of the 9 modules in the female frontloader alcohol drinker network. Only one brain region (the nucleus raphe pontis) was a connector hub in both sexes, suggesting that frontloading in males and females may be driven by different brain regions. In conclusion, alcohol consumption led to fewer, but more densely connected, groups of brain regions in males but not females, and recruited different hub brain regions between the sexes. These results suggest that alcohol frontloading leads to a reduction in network efficiency in male mice.

## Introduction

Frontloading is an alcohol drinking pattern where intake is skewed toward the onset of access which results in intoxication (1). As discussed in our recent review (1), we theorized that alcohol frontloading is driven by the rewarding effects of alcohol and, importantly, alcohol frontloading may predict long-term maladaptive alcohol drinking patterns leading to the development of alcohol use disorder (AUD). Therefore, identification of the brain regions which drive frontloading may lead to the detection of novel AUD risk factors and treatment targets. Despite a growing interest in frontloading in the field, there are no studies to date which identify the brain regions that drive alcohol frontloading behavior. The objective of the current project was to leverage whole brain imaging coupled with graph theory analyses to identify which brain regions / networks are recruited during frontloading.

Advancements in tissue clearing approaches coupled with immunolabeling have opened the door to the unbiased assessment of whole brain networks with single-cell resolution, for reviews, see (2, 3). Immunolabeling-enabled imaging of solvent-cleared organs (iDISCO+) (4, 5) with light sheet imaging is increasingly being utilized in the alcohol field (6–9). Immunostaining for Fos protein, from the immediate early gene c-fos, is commonly paired with the iDISCO+ technique to explore brain wide neural activation associated with a behavioral state (4–6, 8–10). A typical analysis approach is to create functional connectivity matrices through calculating a Pearson correlation between Fos+ cell counts of one brain region and every other brain region from subjects within a given treatment group (6–13). Prior work using these techniques has shown an increase in the strength of positive correlation between most brain regions in alcohol dependent male mice, with a cluster of amygdala regions displaying anticorrelations with the rest of the brain, suggesting that amygdala brain regions may be uniquely involved in the development of alcohol dependence (8). Alcohol drinking has similarly been shown to increase co-activation in the isocortex, cortical subplate, striatum, and pallidum in male mice following four weeks of intermittent access to two-bottle choice drinking as compared to water drinkers (6). However, this effect was opposite in females, with female alcohol drinking mice displaying more negative correlations in most brain regions (6). Together, these results suggest that alcohol drinking results in sex-specific effects on brain region co-activation.

Covariance in activity patterns has been interpreted to reflect groups of brain regions that work together to generate some aspect of behavior. These groups have been characterized as “modules”, which are defined by brain regions that share several, dense connections within the module and (typically) have sparse connections to other brain regions and/or modules (14). It is hypothesized that brain networks are organized in a modular fashion (15) that are subject to remodeling depending on experience or behavioral state (8, 16). Prior work has demonstrated that alcohol dependent mice exhibit a decrease in modularity following alcohol exposure but was not measured under the influence of alcohol. This suggests that alcohol dependence remodels brain networks into a baseline state where they are more correlated (8). Further, this suggests that alcohol dependence results in a less efficient network structure, where more brain regions are recruited to control behavior (8). Other recent work has shown that alcohol re-access following withdrawal leads to increased modularity (7, 9), suggesting that alcohol re-access results in remodeling of brain networks to become more efficient. Additionally, or alternatively, alcohol re-access may serve as a behavioral event capable of recruiting different brain regions as compared to alcohol drinking alone without a withdrawal period (7, 9). Together, these studies indicate that the structure of brain networks is altered by the stage of alcohol abuse, which may reflect differences in how efficiently behavior is generated.

An advantage of using iDISCO+ is the *unbiased* identification of brain regions and modules recruited by alcohol frontloading. This allowed us to determine how alcohol drinking and frontloading lead to remodeling of brain-wide networks. Few studies have assessed alcohol’s impact on the whole brain using immunolabeling and none have assessed alcohol frontloading, thus motivating this approach in the current paper.

Only one published study within the alcohol field has used iDISCO+ whole brain clearing coupled with Fos immunohistochemistry in both sexes, which reported sex differences in co-activation and hub brain regions following 4 weeks of drinking (6). Across strains and species, female rodents typically outdrink males in a variety of alcohol self-administration protocols (17–21); and a recent study provides direct evidence for linking frontloading in female rats as a reason for their greater alcohol intake during operant alcohol self-administration as compared to males (22). Therefore, the current study included both sexes to assess how frontloading and alcohol drinking may differentially impact functional network architecture.

## Methods

Additional detail for some sections is provided in the Extended Methods within the Supplementary Material.

### Subjects

64 adult C57BL/6J mice (32 female, 32 male) were ordered from Jackson laboratories. Mice arrived at the School of Science Vivarium at Indiana University-Purdue University Indianapolis (IUPUI) and were single-housed in standard shoebox cages in a room with a 12-hour reverse light-dark cycle for 7-9 days prior to the beginning of drinking-in-the-dark (DID). Mice were post-natal day 84 ± 3 days on the first day of DID. One male mouse was excluded from all analyses due to a leaky sipper. All procedures were approved by the IUPUI School of Science Animal Care and Use Committee and conformed to the Guidelines for the Care and Use of Mammals in Neuroscience and Behavioral Research (23).

### Volumetric Drinking Monitor Drinking-in-the-Dark

A binge drinking protocol, “drinking-in-the-dark” (DID), was utilized, where mice were given access to 20% EtOH or water (control) for 2 hours a day, 3 hours into the dark cycle for one week (24). On day 8, access was given to the assigned DID fluid for 80 minutes.

During the DID sessions, EtOH or water was consumed from volumetric drinking monitors (VDMs) (Columbus Instruments Inc., Columbus, Ohio, United States) which recorded intake in one-minute bins.

### Blood Ethanol Concentrations (BECs)

50 uL of retro-orbital sinus blood was drawn from all mice on day 8 of DID 80 minutes after DID sipper access. Blood samples were centrifuged in a chilled 4°C centrifuge at 14,000 rotations per minute for 5 minutes. Plasma was then withdrawn and stored at -20°C. BECs were determined using an Analox EtOH Analyzer (Analox Instruments, Lunenburg, Massachusetts, United States).

### Perfusions

80 minutes into drinking on day 8, mice were deeply anesthetized with urethane (injection volume of 0.15 mL with a concentration of 1.5 g/mL) for transcardial perfusion. Frontloaders had an average change point at 20 minutes, *Figures 1B,F*. It is reported that fos expression peaks 60 minutes after behavior (25). Therefore, an average 20-minute change point plus allowing 60 minutes for fos expression to take place provided rationale for the 80-minute perfusion and blood draw time point on day 8. Following perfusion, brains were extracted and placed in paraformaldehyde overnight. The following day, brains underwent 3 30-minute washes shaking in 1x phosphate-buffered saline (PBS). Following the washes, brains were placed in 1x PBS with 0.02% sodium azide at 4°C for up to one week and then were processed according to the iDISCO+ protocol.

**Figure 1.**
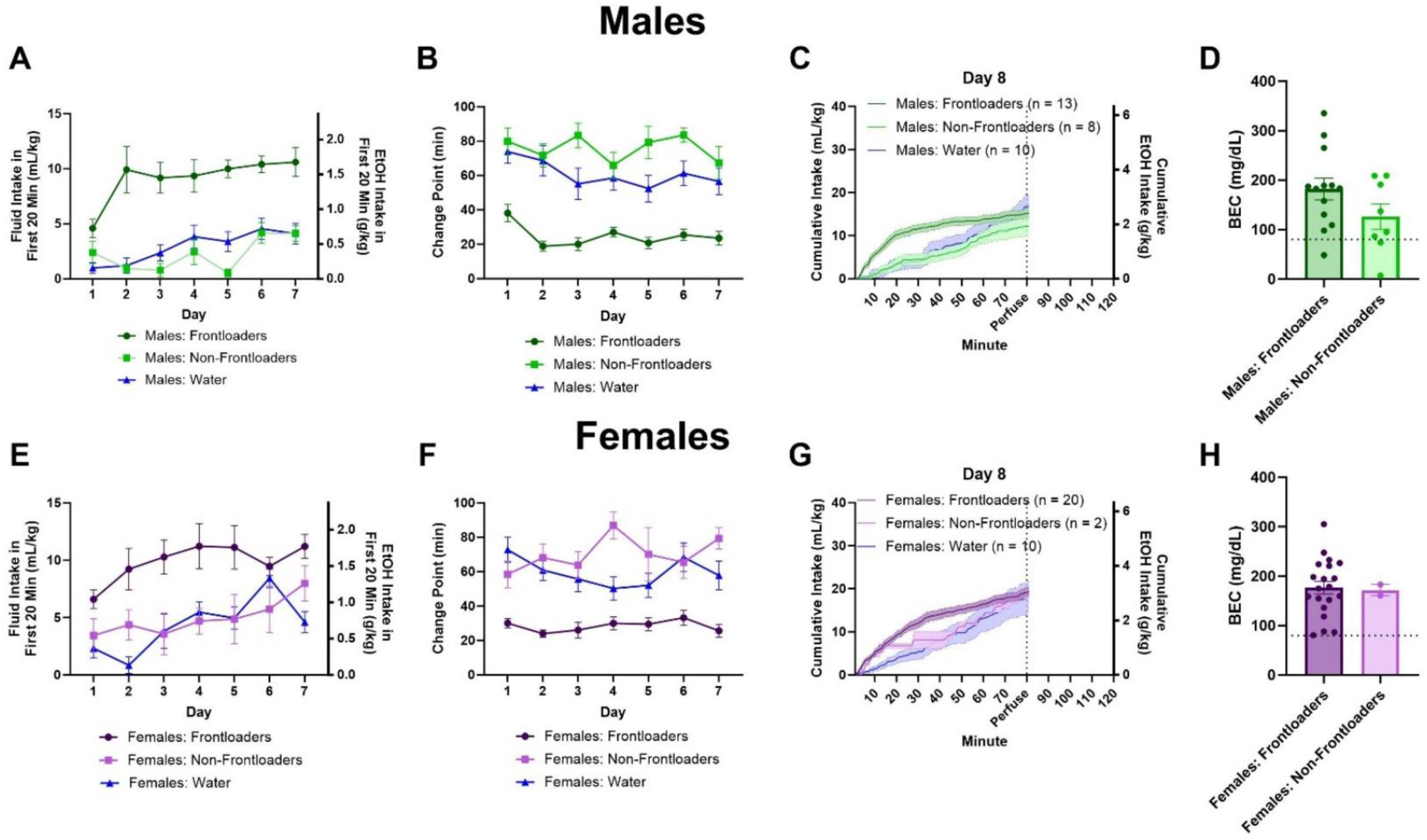
Intakes in the first 20 minutes indicate that male (A) and female (E) frontloaders drink more in the early part of the session. Change point over days indicates earlier change points for both male (B) and female (F) frontloaders, suggesting they consume at a quicker rate than the other groups. Intake patterns on day 8 for males (C) and females (G) are displayed. There were no differences between male (D) and female (H) frontloaders and non-frontloaders in BEC when mice were sacrificed at 80 minutes on day 8. The dashed line on figures D and H represents 80 mg/dL, which is the NIAAA-defined threshold for binge drinking.

### iDISCO+ Tissue Clearing with Fos Immunostaining

iDISCO+ with Fos immunohistochemistry was performed using the protocol reported by the creators of the technique (4, 5). Of note, c-fos rabbit primary antibody (Synaptic Systems, #226 008) was used at a concentration of 1:3000, in PBS with 0.2% Tween and 10 μg/mL heparin/5% DMSO/3% donkey serum at 37°C. The secondary antibody (Donkey anti-rabbit Alexa647, Thermo-Fisher, # A-31573) was used at a concentration of 1:500 in 3% NDS in PtwH at 37°C.

### Light Sheet Imaging

Light sheet imaging was conducted using previously published methods (10). For full detail, please see the Extended Methods in the Supplementary Material.

## Data Analysis

### Statistical Assessment of Frontloading

Mice were categorized daily as frontloaders or non-frontloaders using a change-point detection approach recently described by our laboratory (26). MATLAB code for this analysis is available on Github: https://github.com/cardinger/Detect_Frontloading. Day 8 classification (i.e. frontloader, non-frontloader, or water control group) was used for subsequent brain network analyses as this is the day brains were extracted and the behavior was displayed and captured in the fos expression.

### Statistical Assessment of Behavioral Data

Behavioral data referring to the following were all analyzed: mean daily DID total intake, daily DID intake within the first 20 minutes, and change point. As a main goal of the study was to determine differences between frontloaders and non-frontloaders, these dependent variables were first analyzed using 2 (sex) x 2 (group: frontloaders vs. non-frontloaders) x 7 (day) 3-way mixed methods ANOVAs, first excluding the water group from analyses to determine if there were main or interaction effects of sex within the alcohol groups. Results are in *Supplementary Table 2*. The water group was then added to intake analyses when they were recalculated separately for each sex; see below.

Although many of the behavioral dependent variables indicated main effects of sex, there were interaction effects of sex observed in the brain network data. Therefore, all graphs and subsequent reported behavioral analyses were recalculated using 3 (group: frontloaders vs. non-frontloaders vs. water) x 7 (day) 2-way mixed methods ANOVAs separately for females and males. Greenhouse-Geisser corrections were applied to these analyses when appropriate.

### Statistical Assessment of BECs

Independent samples t-tests (frontloaders vs. non-frontloaders) were used to compare BEC and intake (g/kg) at 80 minutes on day 8. These were calculated separately for each sex.

### ClearMap2.1 to Detect Fos+ Nuclei

Fos+ cells were detected using ClearMap2.1 (https://github.com/ChristophKirst), the current version of the ClearMap software which has been validated in previous whole brain imaging studies (4, 8–10).

### Correlation Matrices

Functional correlation matrices were calculated from Pearson correlations of Log_10_ Fos+ values between brain regions. These data were then organized anatomically using the Allen Brain Atlas for visualization purposes. Further details are provided in the Extended Methods.

### Hierarchical Clustering

Following previous literature (7, 8, 10, 11, 27), the Fos+ Pearson correlations were used to calculate Euclidean distances between brain region pairs. These were calculated separately within sex and group. R studio was used to hierarchically cluster these Euclidean distance matrices. Modules were determined through cutting these dendrogram trees at 50% height.

### Creation of Networks and Identification of Hub Brain Regions

Key brain regions were identified using a graph theory approach for each group and sex (i.e., 5 networks total were assessed: male frontloaders, male non-frontloaders, male water, female frontloaders, female water). The connectivity metrics within module degree z-score (WMDz) and participation coefficient were calculated as described by Guimerà and Nunes Amaral (2005), but modified for networks with weighted edges using a similar approach as previously published research (8). To describe the networks, modules were given names based on regions with the highest WMDz values. Once intramodule connectivity (WMDz) and intermodule connectivity (participation coefficient) were calculated for each brain region, it was possible to categorize the role of each brain region (e.g., hubs or nodes) in each network to identify brain regions of interest. For full detail about this process, please see the Extended Methods. The distribution of types of nodes and hubs (i.e., network cartography) was assessed statistically between groups using chi-square analyses.

## Results

### Total DID Session Intake

In males and females, there was a main effect of day, F(4.185, 111.6) = 4.518, *p* < 0.01 (males), F(4.853, 135.1) = 8.827, *p* < 0.0001 (females), and a main effect of group, F(2, 37) = 29.65, *p* < 0.0001 (males), F(2, 35) = 25.54, *p* < 0.0001 (females), where water-drinking mice consumed more than alcohol-drinking mice and intakes generally increased over days, *Supplementary Figures 1A, B,* respectively.

### Frontloaders Display Greater Alcohol Intake Early in the Session

In males and females, for intake in the first 20 minutes, there was a main effect of day, F(4.561, 120.9) = 3.794, *p* < 0.01 (males), F(4.365, 122.2) = 3.200, *p* < 0.05 (females), where intakes tended to increase over days, and a main effect of group, F(2, 37) = 65.25, *p* < 0.0001 (males), F(2, 35) = 23.02, *p* < 0.0001 (females), where frontloaders consumed more in this early period than non-frontloaders and water drinkers on most days, *Figures 1A, E*, respectively. For change point, there was a main effect of group in both sexes, F(2, 37) = 115.4, *p* < 0.0001 (males), F(2, 35) = 69.78, *p* < 0.0001 (females), where frontloaders had earlier change points than non-frontloaders and water drinkers on most days, *Figures 1B, 1F,* respectively.

On day 8, there was no difference in total intake between frontloaders and non-frontloaders when mice were sacrificed for perfusion and brain extraction 80-minutes into the DID session, *t*(19) = 1.1137, *p* > 0.05 (males), *t*(20) = 0.1139, *p* > 0.05 (females). There was also no difference in BEC at this time, *t*(19) = 1.167, *p* > 0.05 (males), *Figure 1D*, *t*(20) = 0.1141, *p* > 0.05 (females), *Figure 1H,* and most mice regardless of group (i.e., both frontloaders and non-frontloaders) drank to intoxication as indicated by BECs > 80 mg/dL. Intake patterns on day 8 can be seen in *Figures 1C* and *G*.

### Functional Correlation Matrices

Fos functional correlation matrices for males can be visualized in *Figures 2A-C.* One-way ANOVAs calculated to assess differences in R value within anatomical subdivisions between groups were significant for the cortical plate, [F(2, 1132) = 134.5, *p* < 0.0001], striatum, [F(2, 1179) = 42.26, p < 0.0001], hypothalamus, [F(2, 1282) = 3.561, *p* < 0.05], midbrain, [F(2, 835) = 7.610, *p* < 0.01], and hindbrain, [F(2, 1132) = 134.5, *p* < 0.0001]. Šídák’s multiple comparisons post-hoc tests indicated that frontloaders had lower R values in the cortical plate, midbrain, and hindbrain as compared to both control groups. Male frontloaders displayed higher R values in the striatum and hypothalamus as compared to male water drinkers. These results suggest frontloading alters the strength of functional connectivity differently across some anatomical subdivisions, *Supplementary Figure 2*. Note only significant R values (p < 0.05) were included in the ANOVAs.

**Figure 2.**
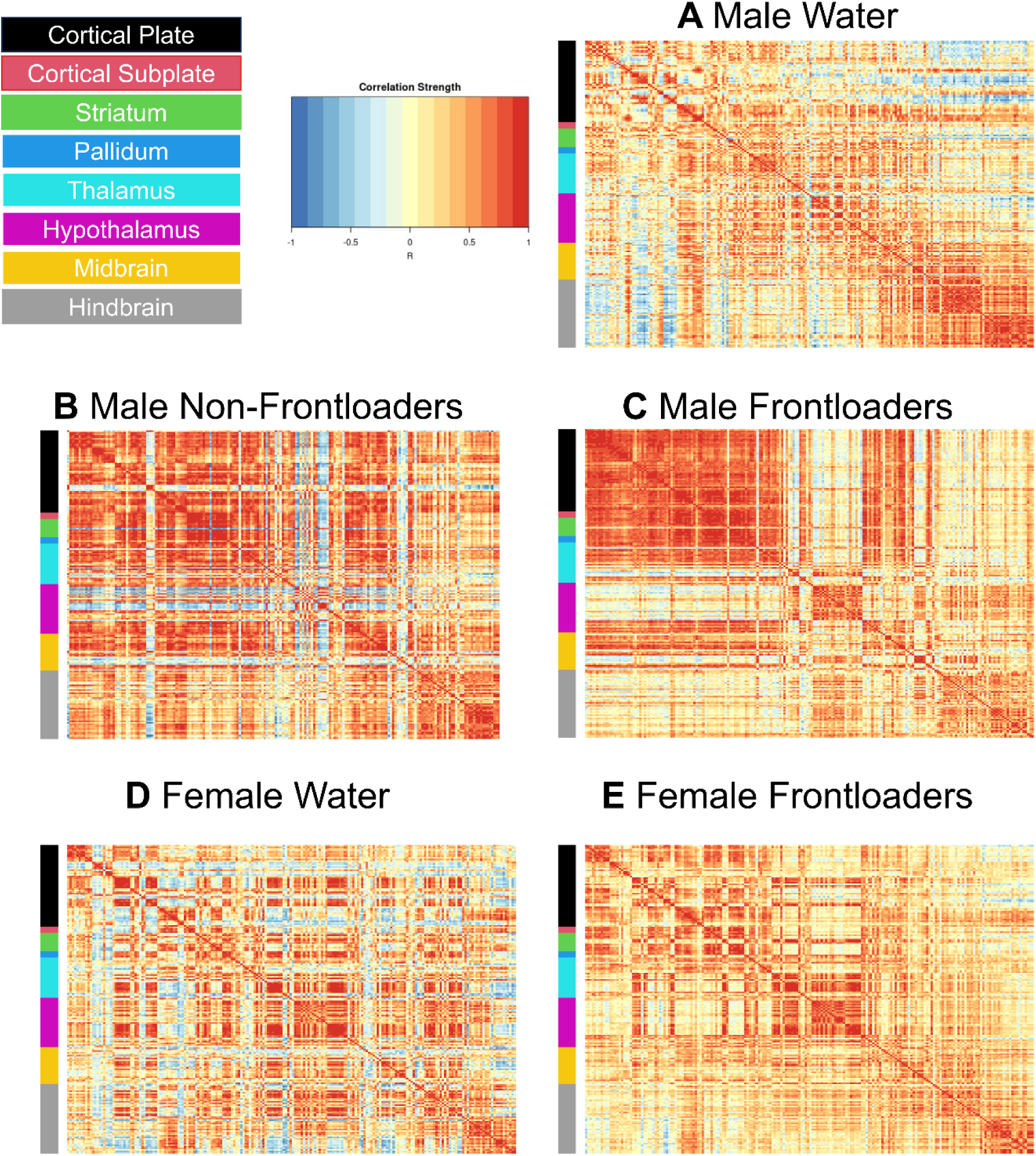
Functional correlation matrices organized using the Allen Brain Atlas. Males (A-C) and females (D, E) are represented. Note that there is no non-frontloading group represented for females as there were only 2 female mice who did not frontload when brains were extracted on day 8, which inhibits the ability to make meaningful conclusions about female non-frontloaders in the current study.

Fos functional correlation matrices for females are shown in *Figures 2D, E.* In order to compare between the sexes, two-way ANOVAs assessing R-values of male and female frontloaders and water drinkers were conducted. Results of this analysis are displayed in Table 1.

**Table 1.** Significant results of the two-way ANOVAs comparing male and female significant R values between brain region anatomical divisions.

|  |  |  |
| --- | --- | --- |
| **Cortical Plate** |  |  |
| *Source of Variation* | *F(DFn, DFd)* | *P value* |
| Sex | F(1, 1558) = 63.99 | p < 0.0001 |
| Group | F(1, 1558) = 73.07 | p < 0.0001 |
| **Cortical Subplate** |  |  |
| *Source of Variation* | *F(DFn, DFd)* | *P value* |
| Sex | F(1, 528) = 27.21 | p < 0.0001 |
| Sex x Group | F(1, 528) = 8.603 | p < 0.001 |
| **Striatum** |  |  |
| *Source of Variation* | *F(DFn, DFd)* | *P value* |
| Sex | F(1, 1457) = 21.76 | p < 0.0001 |
| Group | F(1, 1457) = 50.97 | p < 0.0001 |
| Sex x Group | F(1, 1457) = 156.5 | p < 0.0001 |
| **Pallidum** |  |  |
| *Source of Variation* |  |  |
| Sex x Group | F(1, 568) = 28.35 | p < 0.0001 |
| **Thalamus** |  |  |
| *Source of Variation* | *F(DFn, DFd)* | *P value* |
| Sex | F(1, 2564) = 58.93 | p < 0.0001 |
| Sex x Group |  |  |
| **Hypothalamus** |  |  |
| *Source of Variation* | *F(DFn, DFd)* | *P value* |
| Sex | F(1, 2614) = 6.978 | p < 0.001 |
| Group | F(1, 2614) = 10.61 | p < 0.001 |
| Sex x Group | F(1, 2614) = 81.00 | p < 0.001 |
| **Midbrain** |  |  |
| *Source of Variation* | *F(DFn, DFd)* | *P value* |
| Sex | F(1, 1812) = 189.9 | p < 0.0001 |
| Group | F(1, 1812) = 30.23 | p < 0.0001 |
| Sex x Group | F(1, 1812) = 66.79 | p < 0.0001 |
| **Hindbrain** |  |  |
| *Source of Variation* | *F(DFn, DFd)* | *P value* |
| Sex | F(1, 1558) = 63.99 | p < 0.0001 |
| Group | F(1, 1558) = 73.07 | p < 0.0001 |

Šídák’s multiple comparisons post-hoc tests indicate male frontloaders show higher R values than female frontloaders in all anatomical divisions, *Figure 3.* These results suggest males may be uniquely vulnerable to changes in functional connectivity during alcohol frontloading. Interesting bidirectional interaction effects were noted in the striatum (male frontloaders R value was higher than male water, female frontloaders R value was lower than female water), with an opposite effect in the midbrain.

**Figure 3.**
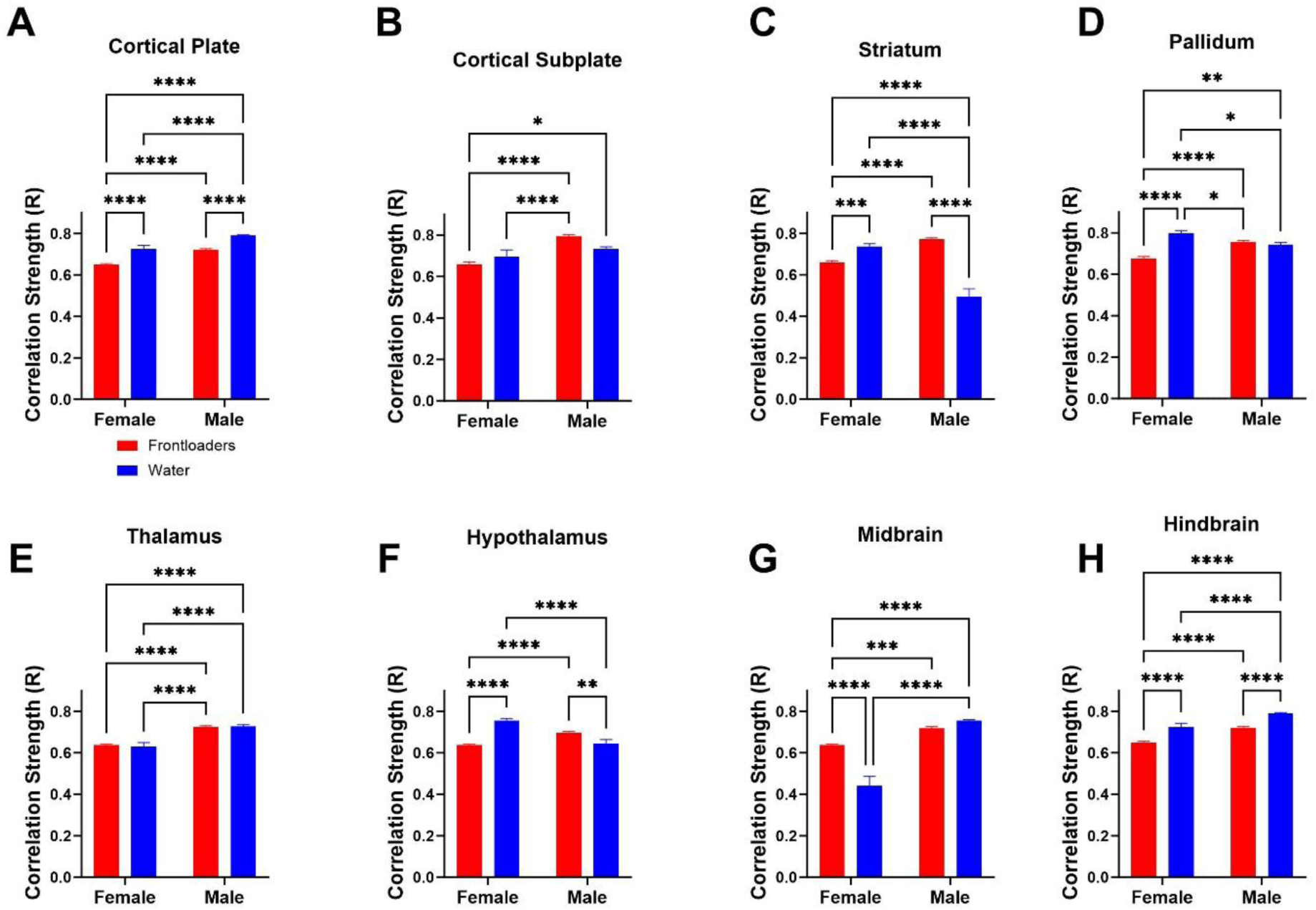
Correlation strength (R value) is compared across groups and sex within anatomical subdivisions. In most anatomical divisions, the strength of correlation is decreased in female frontloaders as compared to water drinkers. This was true in all divisions except for the cortical subplate, thalamus (no differences), and midbrain (where frontloading increased co-activation). Further, there were main effects of sex in all divisions except for the pallidum, with interactions of sex indicated in many divisions. These results suggest that correlation strength is altered differently between sexes, with male frontloaders showing higher R values in all anatomical divisions than female frontloaders.

### Binge Drinking Reorganizes Functional Connections in The Brain

In males, the hierarchical clustering procedure identified 7 modules in water drinkers, *Figure 4A*, 3 in non-frontloaders, *Figure 4B*, and 3 in frontloaders, *Figure 4C*. This decreased modularity identified in alcohol-drinking male mice was observed regardless of the tree-cut threshold used, *Figure 4D*. The modules that each brain region were clustered into are listed in Supplementary Tables 4 (water), 5 (non-frontloaders), and 6 (frontloaders). These results indicate that alcohol binge drinking in males alters the structure of a network.

**Figure 4.**
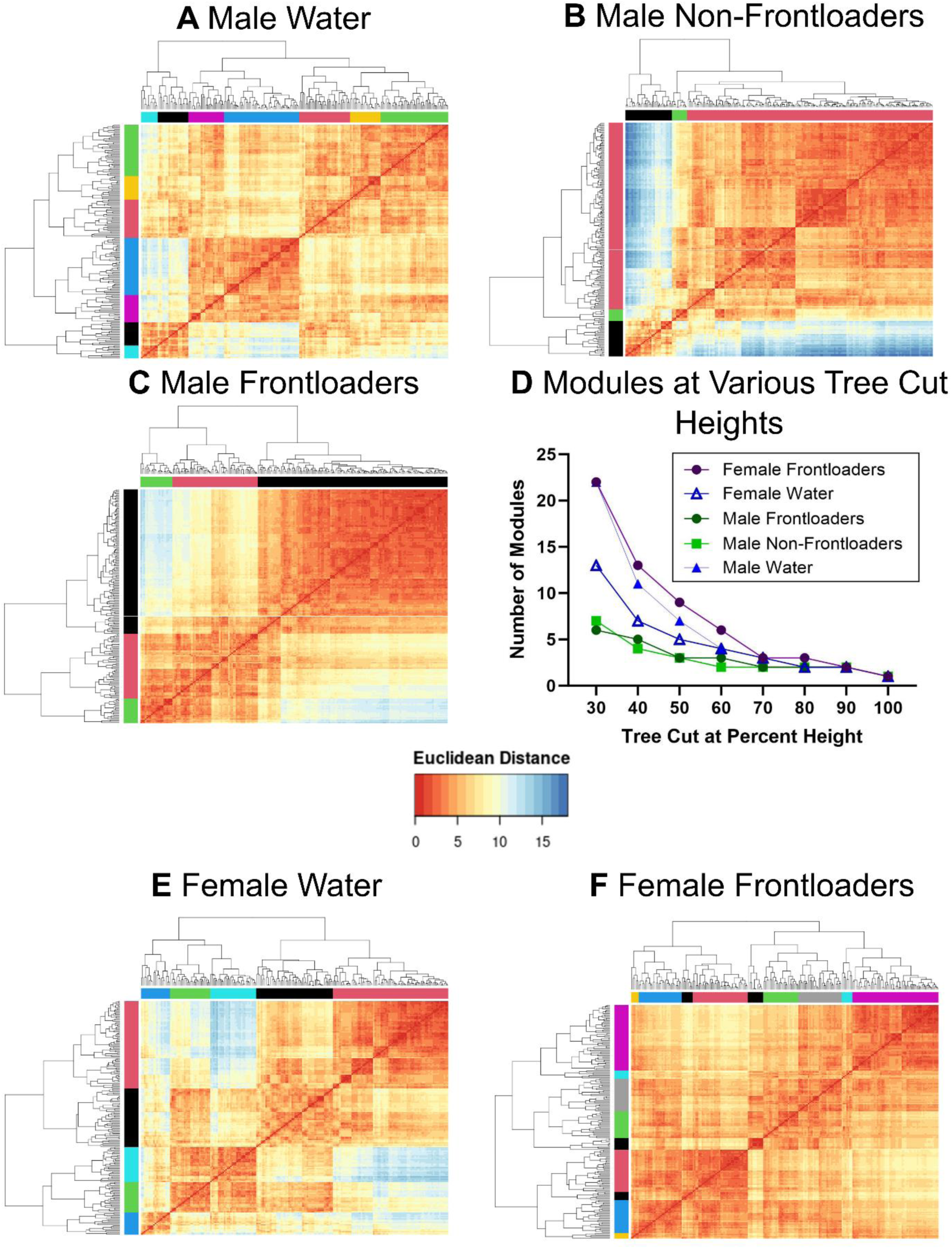
Hierarchical clustering of Euclidean distance matrices for each group. Modules for all groups were determined using a tree-cut height of 50%. In male water drinkers, 7 modules were identified (A). In male non-frontloaders, 3 modules were identified (B). In male frontloaders, 3 modules were identified (C). The decreased modularity in male alcohol drinking groups (i.e. non-frontloaders and frontloaders) is observed regardless of tree cut percentage height chosen (with the exception of extreme cut off values), (D). In female water drinkers, 5 modules were identified (D). In female frontloaders, 9 modules were identified (E). The increased modularity in frontloaders is observed regardless of tree cut percentage height chosen (with the exception of extreme cut off values), (D).

In females, the hierarchical clustering procedure identified 5 modules in water drinkers, *Figure 4E*, and 9 in frontloaders, *Figure 4F*. This increased modularity identified in frontloaders was observed regardless of the tree-cut threshold used, *Figure 4D*. These results suggest alcohol frontloading alters the structure of the network (increased modularity), but interestingly in the opposite way than it did in males.

### Identification of Specialized Nodes in Brain Networks During Alcohol Frontloading

An example of how the relationship between participation coefficient and WMDz was used to characterize every brain region’s role in its network is shown in *Supplementary Figure 3*. Brain regions classified as non-hub connector nodes (having a high participation coefficient), provincial hubs (having a high WMDz), and connector hubs (having both high WMDz and high participation coefficient), were potential key brain regions within a network – each with their own necessary role in the network. Non-hub connector nodes play a role in connecting modules to one another and facilitating connectivity between modules (28). Provincial hubs play a crucial role in how the modular structure of a network is determined and facilitate connectivity within a module (29). Connector hubs enable the flow of information both between and within modules (29). The use of these connectivity metrics to assess prominent network features has proven fruitful in the field of addiction (8, 10). To describe the networks, modules were given names based on regions with the highest WMDz values. Lists of full brain region names and corresponding abbreviations are in *Supplementary Table 3*.

### Male Water Drinkers

Seven modules were identified in the male water drinking functional connectivity network using the hierarchical clustering procedure. There were 1808 edges within this network. One brain region (the VMH) did not have any connections above the 0.7 R value threshold as was thus excluded from this network. Modules in the water drinking network were relatively even in size and can be visualized in *Figure 5.* Further, a relatively even number of connector hubs was observed in each module. A full list of WMDz values, participation coefficient values, module, and categorization for each brain region in the network can be seen in *Supplementary Table 4*.

**Figure 5.**
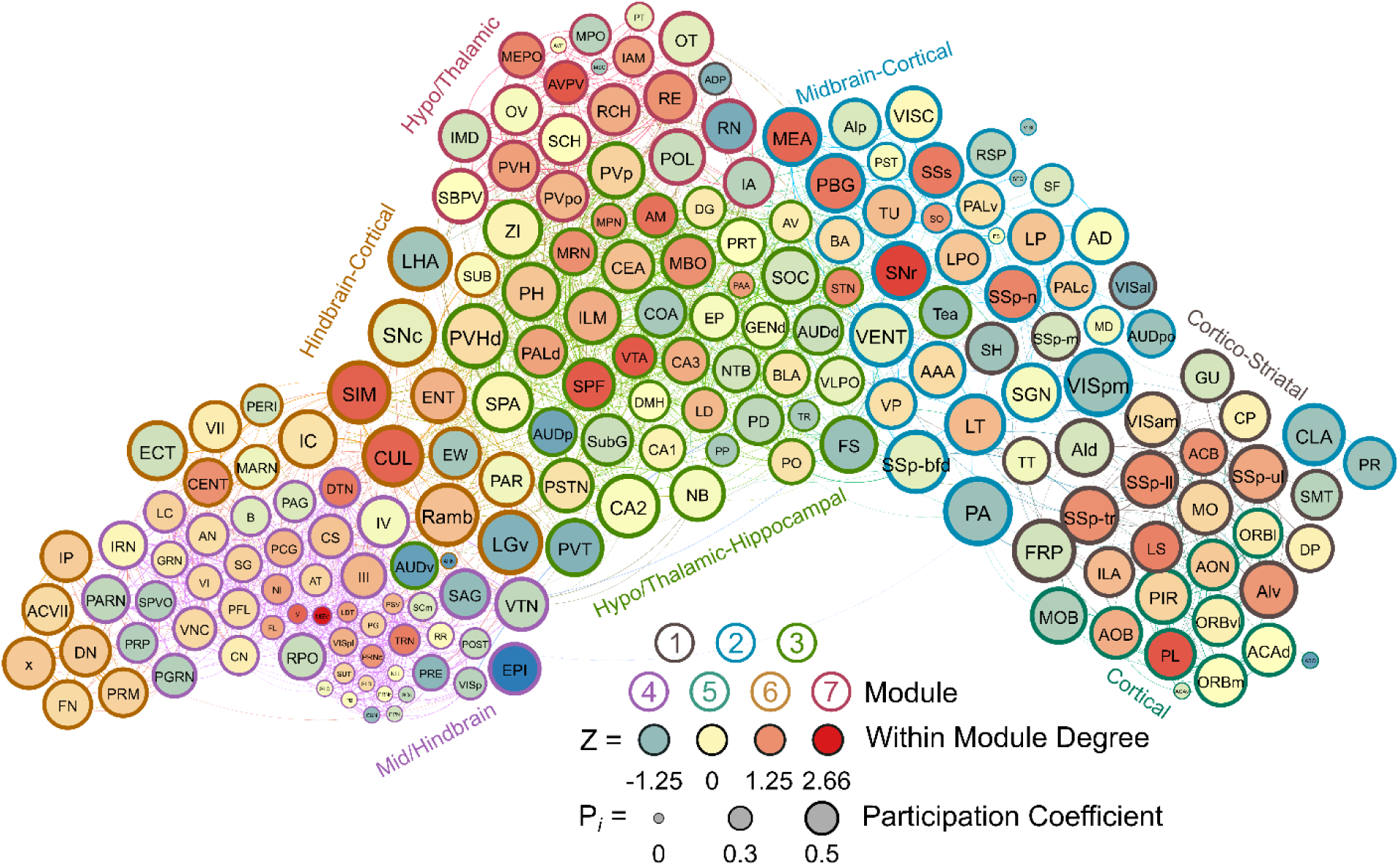
A visualized network of functional connectivity in male water drinkers. Each brain region is a circle. The size of the circle represents the participation coefficient. The color inside the circle represents the WMDz. The color on the outside of the circle represents the module. Seven distinct modules were identified using hierarchical clustering (Figure 4A) and each is represented by a different color.

### Male Non-Frontloaders

Three modules were identified using the hierarchical clustering procedure in male non-frontloaders. There were 4875 edges within this network. All brain regions had at least one connection to another brain region at the 0.7 R value threshold in this network. Module 1 consisted of provincial hubs, peripheral nodes, and ultra-peripheral nodes. This module was driven by intramodular connectivity from hypothalamic-thalamic provincial hubs (e.g. RE, PH, PVpo, SCH, SPA). The peripheral nodes provided sparse connection to the other two modules. In module 1, the PAG connected to the MSC, providing the only direct connection between modules 1 and 2. The CS in module 1 connected to the PG in module 3 and the RPO (module 1) also provided connection to the PG, SUT, and PSV (module 3). Module 2 was the largest module in the network (159 brain regions) and consisted primarily of ultra-peripheral nodes (53%), followed by provincial hubs (31%), and peripheral nodes (16%). Though the cartography of module 2 in male non-frontloaders was similar to that of module 1 in male frontloaders, there were different regions acting as provincial hubs driving within-module connectivity in this largest non-frontloader module. These provincial hubs were comprised of midbrain-hypothalamic-amygdalar-hippocampal regions (e.g. CUN, ZI, CEA, CA3, CA2). The third module was the smallest and consisted only of connector hubs and peripheral nodes. The connector hubs in this region were all from the hindbrain (IP, SPVO, PRM). Interestingly, the peripheral nodes in this module were also all from the hindbrain (e.g. PSV, SUT, x, PG, TRN). This male non-frontloader network is visualized in *Figure 6.* See also *Supplementary Table 5*.

**Figure 6.**
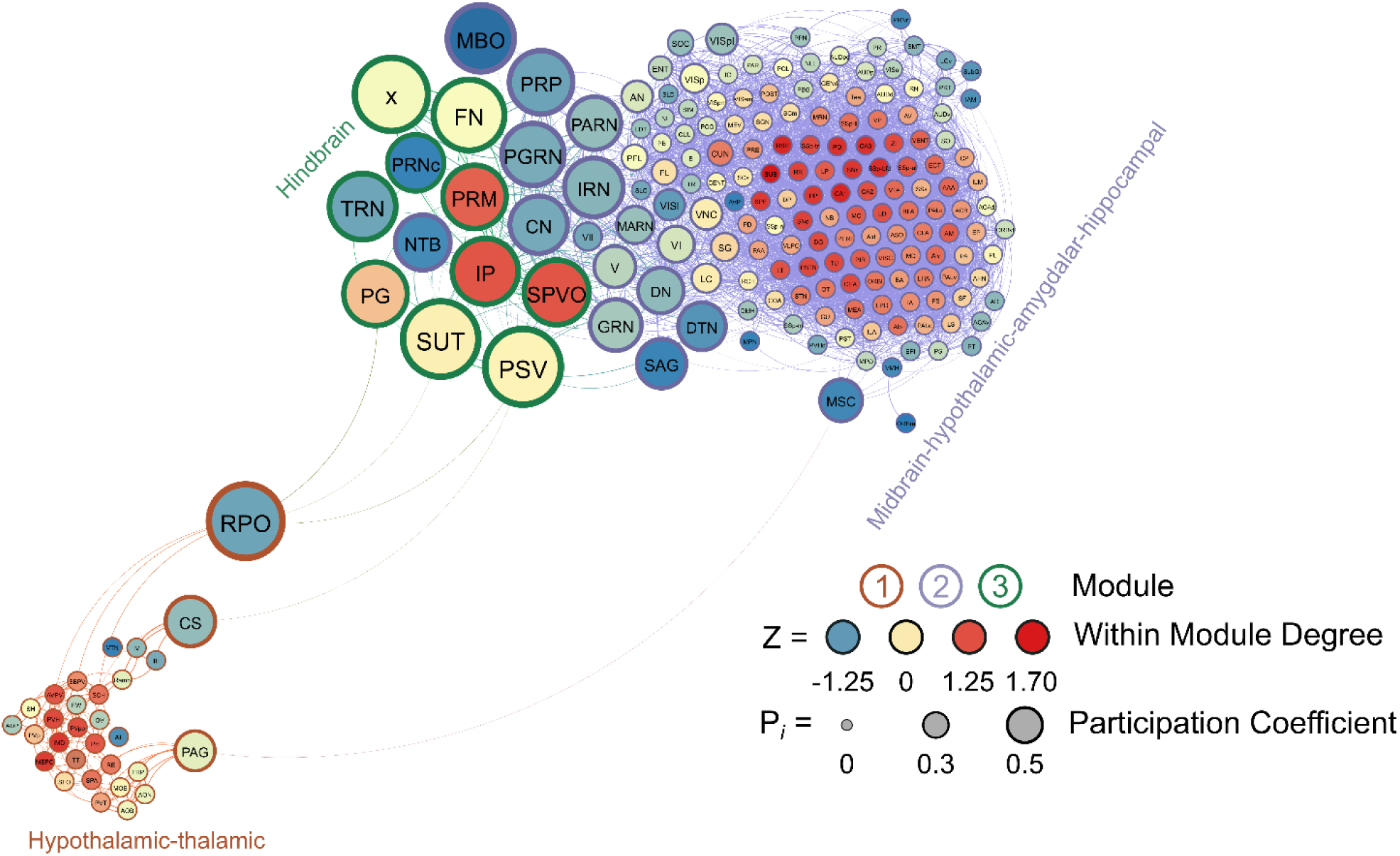
A visualized network of functional connectivity in male non-frontloaders.

### Male Frontloaders

Like in male non-frontloaders, three modules were identified using the hierarchical clustering procedure in male frontloaders. There were 3853 edges within this network. All brain regions had at least one connection to another brain region at the 0.7 R value threshold in this network. Module 1 represented a majority of the network (123 brain regions) and was comprised primarily of ultra-peripheral nodes (61%), then provincial hubs (30%) and peripheral nodes (9%). This largest module was driven by high intramodule connectivity from cortico-striatal-hippocampal provincial hubs, (e.g., CP, Alv, CA3, AAA, LS, ACB, PIR). The peripheral nodes (e.g. VTA, CN, V) provided connection to module 2, but not to module 3. Module 2 was the mid-sized module, and it was driven by hypothalamic-hindbrain connector hubs (e.g., PG, SBPV, RPO, MEPO, PVHd) and provincial hubs (e.g. FL, SCH, LC, IRN). Module 2 also housed the only two non-hub connector nodes in the entire male frontloader network, the MPO and PS (both hypothalamus), suggesting that the intermodule connections facilitated by these hypothalamic-hindbrain connector hubs and non-hub connector nodes in module 2 are crucial for information flow throughout the entire network. This is important as module 1 did not have any direct links to module 3. Module 3 was the smallest module and was driven by hypothalamic-thalamic-mid/hindbrain connector hubs (PVH, RE, and TRN) and a provincial hub (PVT). Of note, the PVT (paraventricular nucleus of the thalamus) had the third largest WMDz within the entire network (WMDz = 1.63). The PVT uniquely stood as a single provincial hub in module 3 (where module 1 had 37 and module 2 had 10 provincial hubs), suggesting that the PVT is uniquely responsible for module 3’s partition within the network and that the PVT plays a large role in intramodule connectivity in this hypo/thalamic-mid/hindbrain module. This male frontloader network is visualized in *Figure 7.* See also *Supplementary Table 6*.

**Figure 7.**
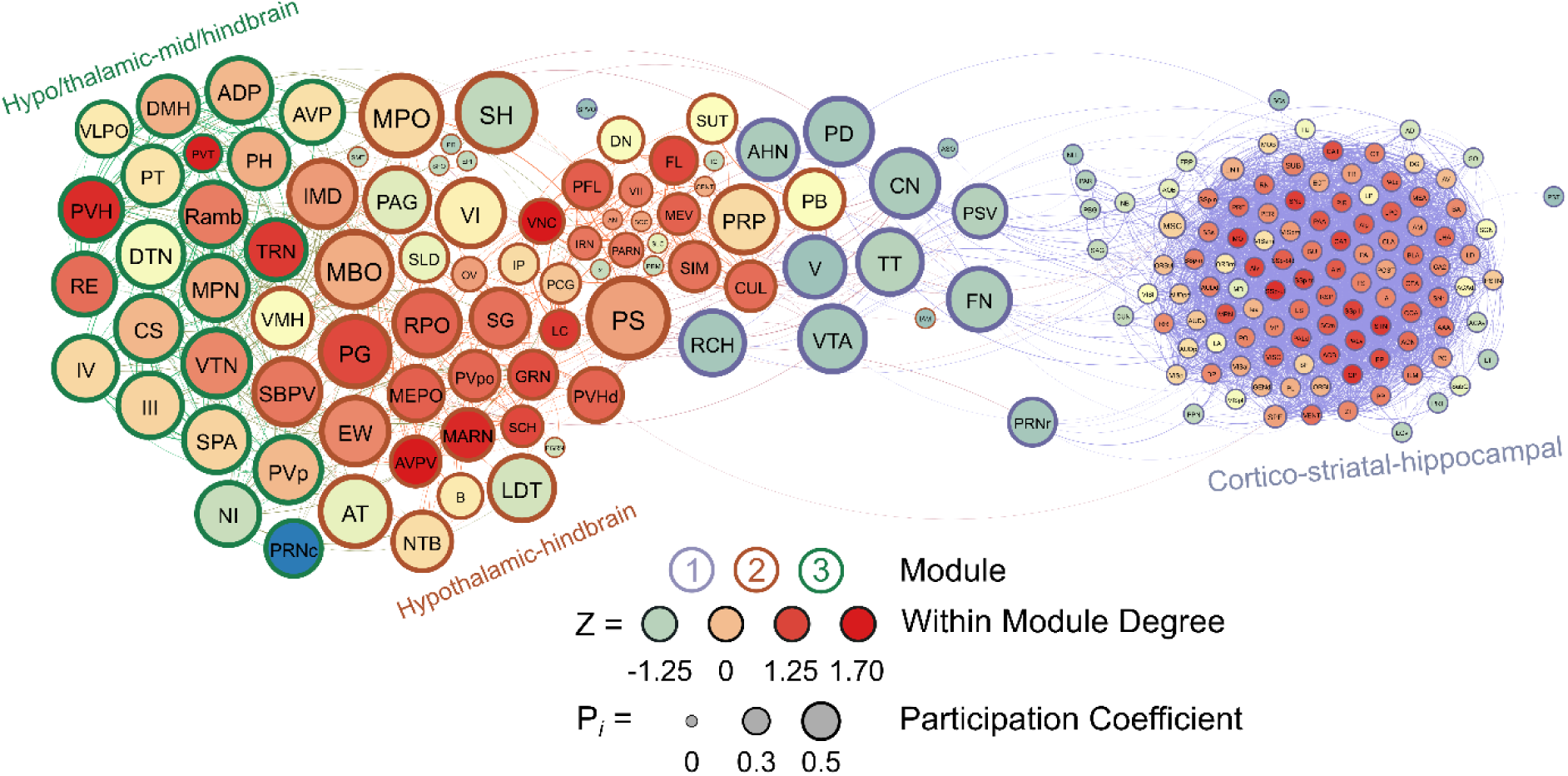
A visualized network of functional connectivity in male frontloaders.

### Female Water Drinkers

Five modules were identified in female water drinkers using the hierarchical clustering procedure. There were 2695 edges in this network. Four brain regions (PP, ASO, ZI, and SAG) did not have any connections above the 0.7 R value threshold as were thus excluded from this network. Interestingly, unlike in the male water drinking nework, not all modules contained connector hubs. Female water drinking modules were driven by connector or provincial hubs, with only 2 of 5 modules containing both types of hubs. Module 1 contained 12 connector hubs and 2 provincial, and both types of hubs were primarily hypothalmic-hindbrain regions (e.g., PRP, SBPV, SG, MEPO). Module 2 was driven by mid/hindbrain-hypothalamic provincial hubs (e.g., B, VTA, DMH, PAG, VLPO). Module 3 was driven by cortico-amygdalar-hindbrain connector hubs (e.g., EP, MEA, V, VII, Alp). Module 4 was driven by visual-auditory provincial hubs (e.g., VISl, VISal, AUDv, AUDp). Lastly, module 5 was driven by thalamo-cortical-hindbrain connector hubs (e.g., PO, CUL, ENT, PFL) and a cortical provincial hub (DG). Module 4 was the smallest, containing only 16 brain regions. The other four modules were relatively even in size. These results suggest that, in female water drinkers, some modules specialize in intermodular communication (those driven by primarily connector hubs, i.e., modules 1, 3, and 5) and some modules specialize in intramodular communication (those driven by provincial hubs, i.e, modules 2 and 4). This female water drinking network is visualized in *Figure 8.* See also *Supplementary Table 7*.

**Figure 8.**
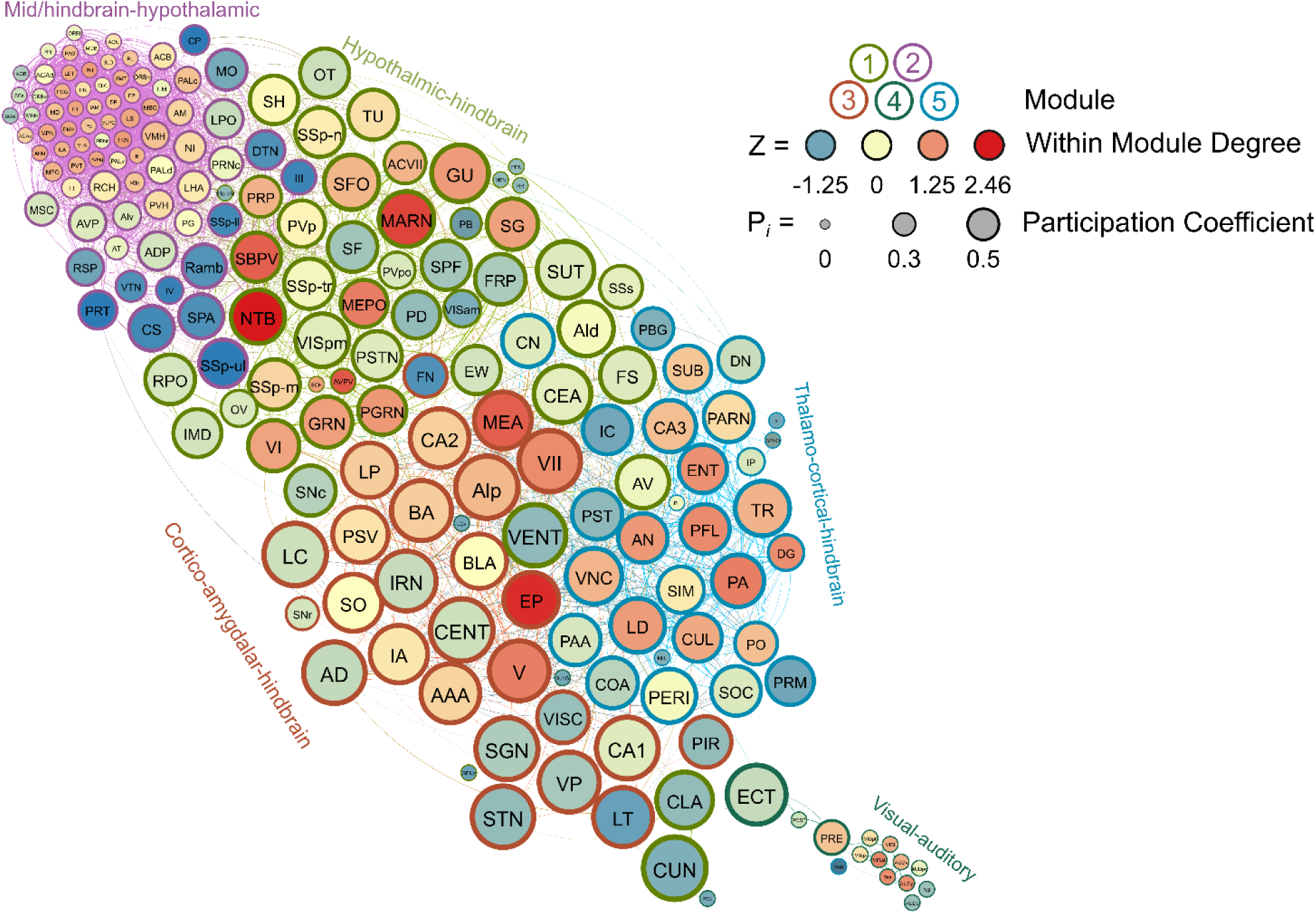
A visualized network of functional connectivity in female water drinkers.

### Female Frontloaders

Nine modules were identified in female frontloaders using the hierarchical clustering procedure. There were 1836 edges in this network. Six brain regions (ASO, CUN, LT, NLL, SOC, VISpm) did not have any connections above the 0.7 R value threshold as were thus excluded from this network. Module 1 was small (7 brain regions) and was driven by the orbital area, lateral part (ORBl) acting as a connector hub. Module 2 was driven by cortico-amydala connector hubs (e.g., PIR, CEA, MEA, BLA). Module 3 had one connector hub, the central lobule (CENT), and 5 provincial hubs which were also predominantly hindbrain regions (e.g., CN, CUL, FL). Module 4 was driven by thalamic-hippocampal connector hubs (e.g., CA3, CA1, LHA, DG). Module 5 was the second smallest module (5 brain regions) and contained one connector hub, the RPO, and one provincial hub, the EW. Module 6 contained no connector hubs, but had 21 provincial hubs which were primarily hypothalamic-thalamic regions (e.g., IAM, LS, AVP, PVHd). Module 7 was the smallest module (4 brain regions) and consisted of one connector hub (AON) and 3 peripheral nodes. This small node primarily consisted of sensory brain regions (AON and MOB for olfaction, LPO for vision), but interestingly also contained nucleus accumbens as a peripheral node. Module 8 contained three connector hubs and 3 provincial hubs from mid-hindbrain regions. Module 9 contained one connector hub and 9 provincial hubs from mid-hindbrain regions. This female frontloading network is visualized in *Figure 9*. See also *Supplementary Table 8*.

**Figure 9.**
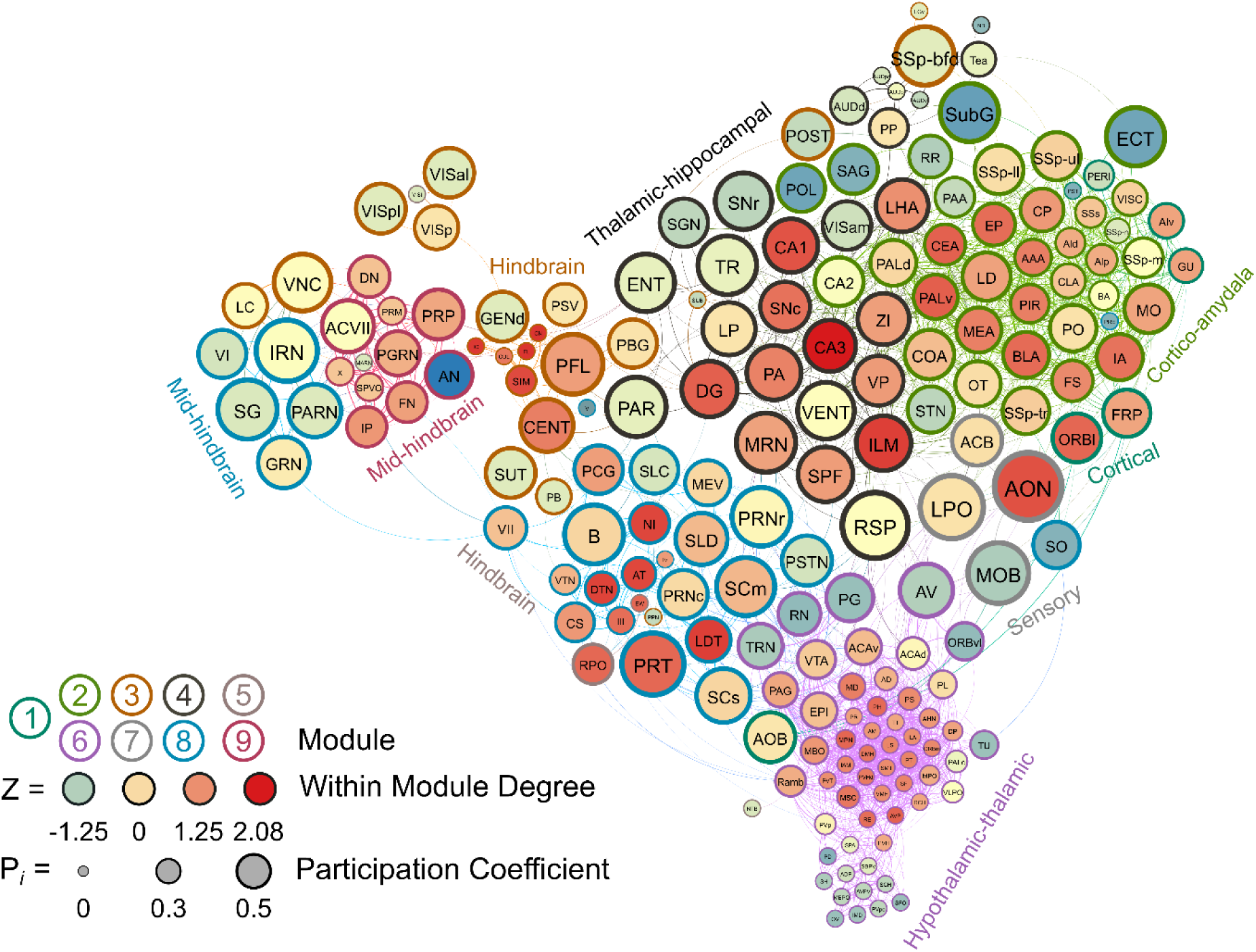
A visualized network of functional connectivity in female frontloaders.

### Binge Alcohol Drinking Alters the Network Cartography of Male, but not Female, Mice

Chi-square analyses indicated that male frontloaders displayed network cartography which differed from the male water group, Χ^2^(4) = 120.4, *p* < 0.0001. Similar results were found when comparing the male non-frontloaders to the male water group, Χ^2^(4) = 178.9, *p* < 0.0001. In contrast, female frontloaders and female water drinkers did not differ, Χ^2^(4) = 2.941, *p* > 0.05. Overall, male water drinkers had more connector hubs (with better distribution throughout modules), more non-hub connector nodes, and fewer ultra-peripheral nodes, which indicated more intermodular communication in water drinkers than non-frontloaders and frontloaders, *Figure 10A-C*. In contrast, female water drinkers and frontloaders had a strikingly similar network cartography, *Figure 10 D,E*.

**Figure 10.**
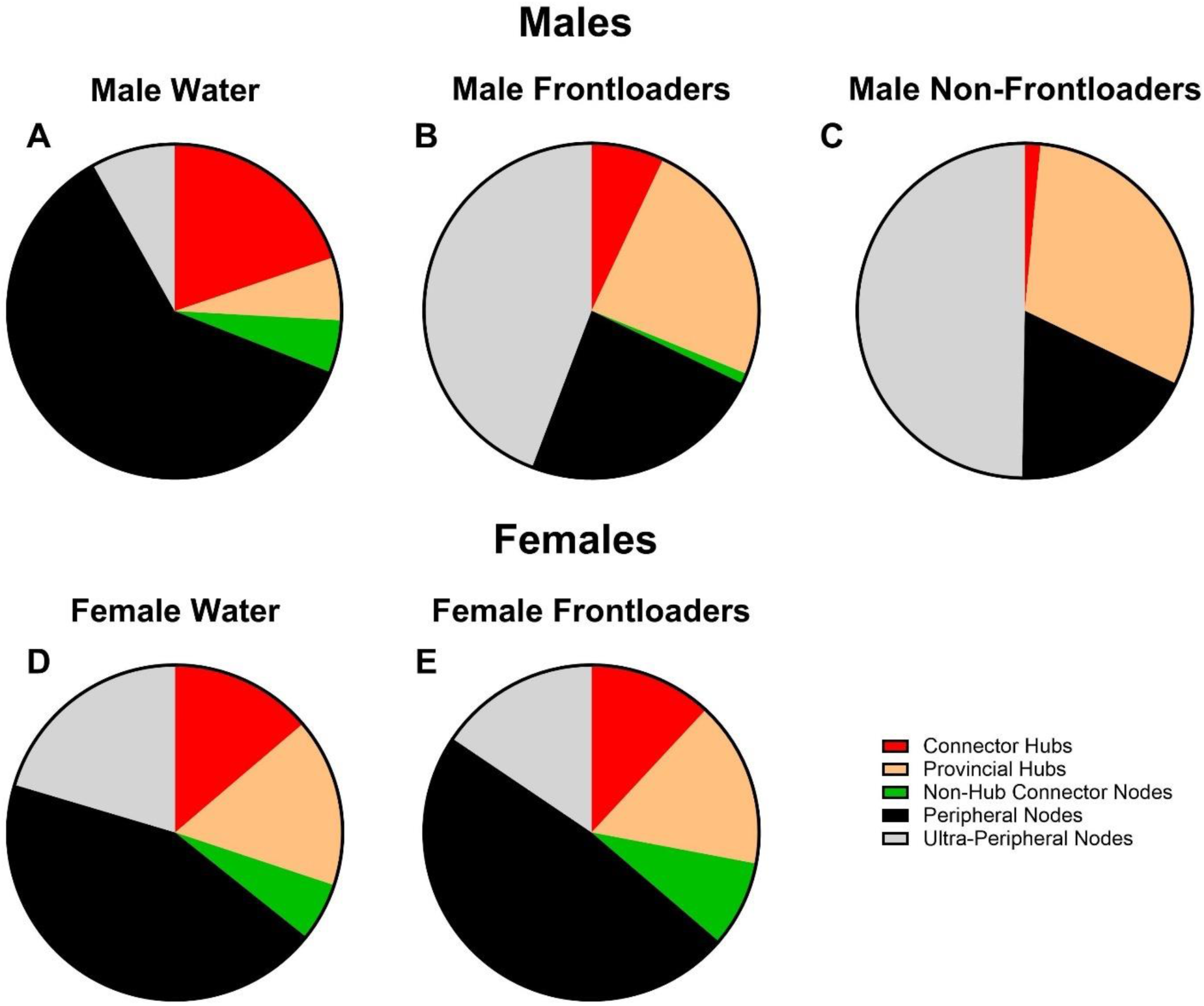
Network cartography. Male water drinkers (A) had more identified connector hubs and non-hub connector nodes - and fewer ultra-peripheral nodes - as compared to male non-frontloaders (B) and male frontloaders (C). These results suggest that water drinking mice have a more globally connected brain network. Female water drinkers (D) and female frontloaders (E) displayed a similar breakdown of types of hubs and nodes within their respective network.

### Brain Regions That May Drive Frontloading

To consider which brain regions may be important in frontloading behavior, lists of connector hubs (high WMDz and high participation coefficient), provincial hubs (high WMDz) and non-hub connector nodes (high participation coefficient) were compared within sex between groups. Brain regions in these categories, with overlap between frontloaders and non-frontloaders or frontloaders and water drinkers, were considered to not be uniquely important in frontloading behavior. A list of connector hubs, provincial hubs, and non-hub connector nodes with no overlap between frontloaders and their within-sex control groups was generated. Graphs of those regions can be seen in *Figures 11A* and *B* for males and females, respectively.

**Figure 11.**
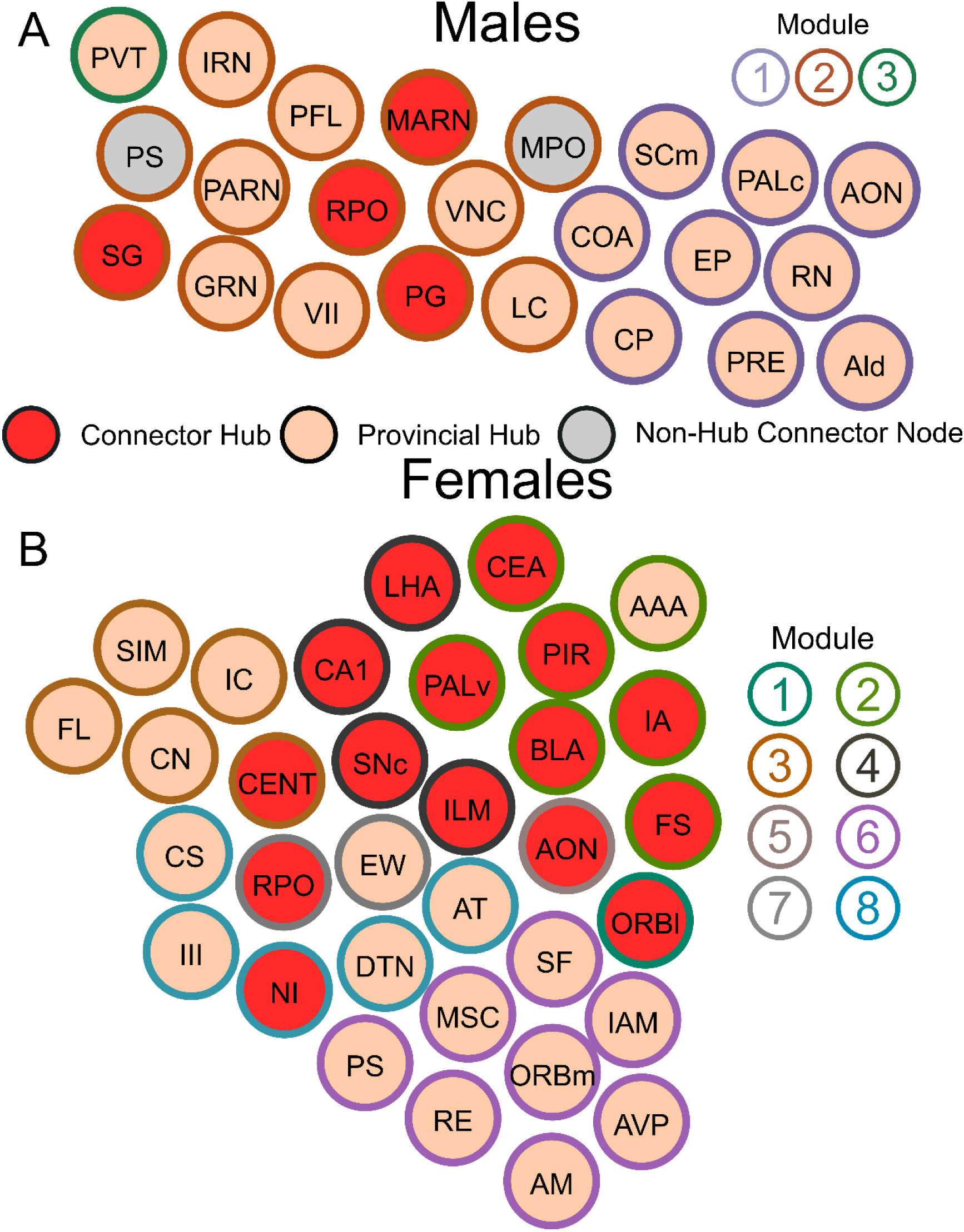
Connector hubs, provincial hubs, and non-hub connector nodes which were unique to male frontloaders (A) and female frontloaders (B). Each brain region is a circle. The color inside the circle represents the role of the brain region in the male frontloader network. The color on the outside of the circle represents the module. As these brain regions were not key brain regions in male non-frontloaders or male water drinkers, they may play a unique role in alcohol frontloading.

## Discussion

In males, alcohol frontloading increased co-activation in the striatum and hypothalamus compared to water drinking. In contrast, frontloading decreased the strength of correlation compared to both non-frontloaders and water drinkers in the cortical plate, midbrain, and hindbrain, suggesting that alcohol frontloading results in recruitment of brain regions divergent from binge drinking which does not use a frontloading pattern, *Figure 2, Supplementary Figure 2*. Alcohol drinking (both frontloading and non-frontloading) decreased modularity and led to the recruitment of more ultra-peripheral nodes and fewer connector hubs as compared to water drinking, *Figures 4, 10*. This difference in network cartography between alcohol drinkers as compared to water drinkers is similar to a recent MEG study which reported a loss of hub regions as non-human primates transitioned from early to chronic heavy drinking (30). Although the same number of modules was identified in male frontloaders and non-frontloaders, these networks displayed some differences in key brain regions involved. Ultimately, four connector hubs and 17 provincial hubs were uniquely identified in male frontloaders (i.e., were brain regions that did not have this status in male non-frontloaders or water drinkers), *Figure 11A*.

In female frontloaders, correlation strength was decreased as compared to water drinkers in all anatomical divisions except for the cortical subplate and thalamus (no differences) and the midbrain (where frontloading increased co-activation as compared to water drinking).

Additionally, it was found that male frontloaders had greater co-activation in all anatomical divisions as compared to female frontloaders, *Figure 3*. A limitation in the current study is the inability to calculate network activity for female non-frontloaders due to a low *n* in this group. Therefore, the conclusions in females cannot be made about alcohol frontloading specifically compared to non-frontloading alcohol drinking but do allow for conclusions about how alcohol binge drinking rearranges functional networks in females. The current results are similar to those of prior studies which have reported a decrease in correlation strength in isocortex, cortical subplate, striatum, and pallidum in female alcohol drinkers as compared to both female water drinkers and male alcohol drinkers (6). Prior work has also shown that female binge drinking mice display lower correlation strength in most brain regions as compared to water controls (7). In the current study, it was found that alcohol binge drinking increased modularity compared to water consumption, which is similar to previous results (7). Lastly, 16 connector and 17 provincial hubs were uniquely identified which were distributed across 8 of the 9 modules in the female alcohol drinker network, *Figure 11B*.

### Sex Differences in Frontloading: Do Males and Females Binge Drink for Different Reasons?

The current study identified differences in functional network activation between the sexes. In males, the unique key brain regions identified in frontloaders (*Figure 11A*) included overlap with brain regions known to play a role in drug seeking and withdrawal (31). The paraventricular nucleus of the thalamus (PVT) was uniquely important in the male frontloading network as it was the sole provincial hub in the hypo/thalamic-mid/hindbrain module, suggesting that the PVT is responsible for this module’s partition within the network. More generally, the PVT has recently been described as playing an integral role in regulating homeostatic behavior in part through incorporating information about prior experience (32). In this regard, the PVT and it’s circuitry may be of interest for future work aimed at manipulating frontloading behavior, as it is established that alcohol frontloading is driven by prior experience with alcohol (26). A hypothesis which arises from the current findings is that male mice engage in alcohol frontloading to reach a desired set point of intoxication which is motivated from learned experiences with alcohol drinking. This set point may be regulated through the PVT, though further research is needed to assess what causal role, if any, the PVT plays in alcohol frontloading.

In females, 16 connector and 17 provincial hubs were uniquely identified (*Figure 11B*). The Edinger-Westphal nucleus (EW) was identified as a provincial hub in the same module as the RPO. The RPO was the only unique connector hub with overlap between both sexes, and it has been shown that urocortin-positive neurons in the EW are important for escalation of alcohol intake (33). Several amygdala brain regions were also uniquely identified as hubs in female frontloaders. These included the BLA and IA as connector hubs, with the AAA acting as a provincial hub. Of note, there was not a prominent group of amygdala regions identified as playing a hub role in male frontloaders. It is well established that the amygdala has a role in alcohol drinking. Both the BLA and CEA show dysregulation of corticotropin releasing factor (CRF) circuitry (34–42). The CEA has known roles in regulating negative affective states, which has been shown to relate alcohol drinking and dependence (43). It is possible that this recruitment of many amygdala regions in female frontloaders suggests a role of negative reinforcement in female binge drinking / frontloading. Our recent review discussed the idea that while negative reinforcement is not necessary for the development of alcohol frontloading, it does exacerbate it (1), also see (44, 45). Therefore, females drinking alcohol for negative reinforcement is a viable hypothesis, but future research is needed to determine the relationship between female binge drinking, frontloading, and negative reinforcement.

An additional or alternative hypothesis that arises from the prominence of amygdala regions in females is that binge drinking is driven by stress or anxiety. In addition to their roles in addiction, the CEA and BLA are validated to regulate responses to anxiety and fear (39, 46–48). Recent work suggests that women are more likely to drink to cope with stress and anxiety as compared to men (49), and women who experience social stress reach higher BECs in a subsequent open access drinking test as compared to men (50). Though the current protocol did not measure cortisol levels or conduct any anxiety assays, the recruitment of many amygdala brain regions as hubs in female alcohol binge drinkers, but not males, raises the hypothesis that female C57BL/6J binge drinking may in part be driven by anxiety. This is an avenue for future research.

## Conclusions

In conclusion, the current study identified sex differences in modularity and brain regions expressing co-activation following binge alcohol drinking. Together, these results indicate that binge alcohol drinking remodels the functional architecture of networks differently between the sexes, leading to fewer, but more densely connected, groups of brain regions in males but not females. These results suggest that alcohol frontloading leads to a reduction in network efficiency in male, but not female, mice. Further, in males, these results suggest that frontloading specifically differs from non-frontloading in terms of which brain regions displayed co-activation, leading to the recruitment of different brain regions as hubs when a frontloading drinking pattern is used.

