## Supplement for "Sex Differences in Neural Networks Recruited by Frontloaded Binge Alcohol Drinking"

### SUPPLEMENTARY MATERIAL

#### Extended Methods

***Subjects***

Cage changes occurred one day prior to the start of DID and were not performed again during the 8-day DID experiment. Mice had *ad libitum* access to LabDiet 5001 (a standard rodent chow), including during DID testing. Mice had *ad libitum* access to water through standard home cage water bottles, except during the DID sessions where their standard home cage water bottles were replaced with volumetric drinking monitor sippers containing 20% EtOH (or water if they were in the water drinking group). Home cage water bottles did not utilize the volumetric drinking sippers but did contain sippers with identical-sized drinking orifices.

***iDISCO+ Tissue Clearing with c-fos Immunostaining***

All washes and incubations were conducted using a shaker. All steps were conducted at room temperature unless otherwise specified. On day 1, brains were pre-treated with increasing concentrations of methanol (MeOH) diluted in MilliQ water. Brains were placed in the MeOH solutions for one hour each: 20%, 40%, 60%, 80%, and then 100% (2 times). Brains were then placed in 33% MeOH/66% dichloromethane (DCM) in a fume hood overnight. On day 2, brains were washed in 100% MeOH and then chilled for 10 minutes at 4˚C. Brains were then bleached with chilled 5% hydrogen peroxide in MeOH overnight at 4˚C. On day 3, brains were rehydrated with decreasing concentrations of MeOH. Brains were placed in the MeOH solutions for one hour each: 80%, 60%, 40%, 20%, then 1x PBS containing 0.02% sodium azide. Brains were then washed twice in PBS/0.2% TritonX-100/0.2% sodium azide for one hour each wash. Brains were then incubated in PBS/0.2% TritonX-100/20% dimethyl sulfoxide (DMSO)/0.3 M glycine/0.02% sodium azide (permeabilization solution) overnight at 37 ˚C. On day 4, brains remained in the permeabilization solution. On days 5 and 6, brains were incubated in PBS/0.2%

TritonX-100/10% DMSO/6% donkey serum/0.02% sodium azide (blocking solution) at 37˚C. On day 7, brains were incubated in c-fos rabbit primary antibody (1:3000, Synaptic Systems, #226 008) in PBS with 0.2% Tween and 10 µg/mL heparin (PTwH)/5% DMSO/3% donkey serum at 37˚C. This primary antibody concentration was selected following a test of 1:1000, 1:3000, and 1:10000 in 6 mice (1 of each sex at each concentration), perfused after 70 minutes of DID on day 8 in experiment 1. Brains remained in the primary antibody for one full week (day 7-the morning of day 14). On the morning of day 14, brains were washed with PtWH 5 times, one hour per wash. Brains remained in the final PtWH wash overnight. On day 15, the secondary antibody was applied. Brains were also moved to opaque black tubes to prevent photobleaching. Donkey anti-rabbit Alexa647 (1:500, Thermo-Fisher, # A-31573) was placed in 3% NDS in PtwH at 37˚C and brains remained in this secondary antibody solution until the morning of day 22. On the morning of day 22, brains were washed with PtWH 5 times, one hour per wash. Brains remained in the final PtWH wash overnight. On day 23, brains were dehydrated with increasing concentrations of MeOH. Brains were placed in the MeOH solutions for one hour each: 20%, 40%, 60%, 80%, and then 100%, then were placed in a fresh solution of 100% MeOH overnight. On day 24, brains were incubated in 66% DCM/33% MeOH for 3 hours, then washed twice (15 minutes per wash) in 100% DCM to remove any remaining MeOH. Brains were then stored in dibenzyl ether in new black opaque tubes until they were imaged. For a list of reagents used and their supplier, see *Supplementary Table 1.*

***Statistical Assessment of Frontloading***

In brief, a change point algorithm was applied to determine 3 change points in the DID session where the rate of intake differed the most in the session. Then three criteria were assessed, where the change point with the best fit statistically was the reference for all criteria. 1) The change point with the best fit statistically was the earliest change point and/or was within the first half of the session. (2) The pre-change point slope was significantly greater than the post change-point slope, as determined through a *t*-test comparing beta weights of pre versus post change point regressions. (3) The pre-change point slope exceeded the rate of alcohol metabolism. If all three criteria were met, subjects would be categorized as frontloaders. If any criteria were not met, subjects were categorized as non-frontloaders.

VDM sipper access was staggered by 5 minutes on all days of DID testing to account for the need to have time to perfuse all mice 80 minutes after their DID sipper access day 8. An additional criterion was applied to the change point frontloading classification analysis where mice had to consume greater than 16.667% of their total intake within the first 20 minutes to be considered frontloaders on day 8 (in addition to meeting all 3 change point criteria described above). It should be noted that 20 minutes represents 16.667% of a two-hour DID session, therefore it was expected that mice would have greater than 16.667% of their total intake within 20 minutes if the mice were displaying frontloading behavior. The 20-minute timepoint is of interest as this is the average change point for frontloaders (Figures 10I, 11I) and was used to determine the 80-minute perfusion timepoint. This ensured that mice in the frontloading group were displaying frontloading behavior within the 20-minute time point of interest to be captured by c-fos activity. Including this intake percentage criterion only reclassified two male mice from the frontloader group to the non-frontloader group on day 8 (as compared to if the change point criteria were used without this additional criterion).. Day 8 classification (i.e. frontloader, non-frontloader, or water control group) was used for subsequent brain network analyses as this is the day the brains were extracted and the behavior was displayed and captured in the c-fos expression.

***Light Sheet Imaging***

Brains were imaged using a light-sheet microscope (UltraMicroscope II, Miltenyi Biotec) equipped with a sCMOS camera (Andor Neo) and zoom body configuration. Imspector Pro software 7.1.4 (Lavision Biotec, Bielefeld, Germany) controlled the microscope. The microscope had a NKT Photonics SuperK EXTREME EXW-12 white light laser with three fixed light sheet generating lenses on each side. Filters at 488 and 640 nm were used to capture autofluorescence and c-fos image stacks, respectively. Right hemisphere scans were completed at 0.8x magnification (1.6x effective magnification) with a light sheet numerical aperture of 0.026. Brains were first scanned for autofluorescence at 488 nm with an exposure time of 50 ms, step-size of 10 um, laser transmission control of 50, and sheet width of 100%. Brains were then scanned for c-fos at 640 nm with an exposure time of 50 ms, step-size of 3 um, laser transmission control of 100, and sheet width of 100. The dynamic focus feature was used when scanning at 640 nm. The imaging resolution was (x = 3.780 μm, y = 3.780 μm, z = 3 μm) for the c-fos 640 nm image stacks and (x = 3.780 μm, y = 3.780 μm, z = 10 μm) for the 488 nm autofluorescence image stacks.

***ClearMap2.1 to Identify fos+ Cells***

Image stacks were individually cropped to be processed from the bottom of the olfactory bulbs to the beginning of the cerebellum. The individual cropping was necessary as not all brains were the exact same size (e.g., female brains tended to be slightly smaller than male brains). The autofluorescence images captured using the 488 nm filter were registered to a reference image stack which has known corresponding brain region divisions from the Allen Brain Adult Mouse 25μm Atlas (4). This registered Allen Brain Atlas map was then used to delineate brain regions for the c-fos image stack captured with the 640 nm filter. Each brain was individually visually inspected for appropriate alignment to the Allen Brain Atlas reference using the hippocampus as a primary landmark.

Previous validation of the ClearMap pipeline indicates 99% accuracy in the automatic detection of fos+ cells as compared to a manual fos+ cell counting approach (4). The cell detection parameters (20 pixel diameter, 3 pixel maxima) used were identical to the well-validated Renier et al. (2016) publication and others (11). However, a more conservative cell filter was applied (min 50 voxels, max 400 voxels) due to overcounting with the default cell filter (min 0 voxels, max 700) in our dataset. These same cell detection and filter parameters were applied to every brain to ensure that differences in fos+ cell counts between groups were due to actual differences between groups and not differences in fos+ cell detection method. Using this ClearMap2.1 pipeline, 441 brain regions with fos+ cells were identified. For easier interpretation of analyses, these brain regions were collapsed into larger divisions. For example, initially, distinct cell counts were identified for the lateral vestibular nucleus, medial vestibular nucleus, spinal vestibular nucleus, and superior vestibular nucleus. Fos+ cell counts from all four of these brain regions were then added together into one brain region named "vestibular nuclei." Brain regions were only collapsed once, e.g., vestibular nuclei were not then also collapsed with the rest of the brain regions which comprise the "medulla, motor related" within their Allen Brain Atlas division. After collapsing brain regions once, 200 brain regions were identified. The number of brain regions identified using this approach can vary based upon how many divisions are collapsed. Previous studies using similar techniques have reported fos+ cells from 178 (11), 123 (8), 306 (6), and 110 (9) regions. As some brain regions are larger than others, all fos+ counts were log10 transformed to facilitate comparison of activation in the whole brain. This approach of using log transformation to normalize whole brain fos+ cell counts is common practice in the field (8, 10, 11, 53, 54).

***Correlation Matrices***

Functional connectivity matrices were calculated separately for each group and sex. Note that only two female mice were in the non-frontloading group on day 8. Therefore, it was not possible to calculate correlation matrices and brain network diagrams for female non-frontloaders. $Log_{10}$ Fos+ values were used to calculate Pearson correlations from every brain region to each of the other regions across mice within a group. These matrices were then organized anatomically using the Allen Brain Atlas with the following divisions annotated: cortical plate, cortical subplate, striatum, pallidum, thalamus, hypothalamus, midbrain, and hindbrain. Note that this organization did not change the correlation values and was used for visualization only. This method of creating functional connectivity matrices using Pearson correlations and then organizing them anatomically is common practice in this field (6-11, 16, 17, 59).

To assess if there were differences between sex and group, R values in each matrix were thresholded at $p < 0.05$ to only include significant correlations for group comparisons. Anatomical divisions (cortical plate, cortical subplate, striatum, pallidum, thalamus, hypothalamus, midbrain, and hindbrain) were analyzed separately using 2 (sex) x 2 (frontloaders vs. water) 2-way ANOVAs. Sidak's multiple comparisons post-hoc tests were then used to further assess significant effects of sex and group. A similar analysis was completed in only males to include their non-frontloading group: a one-way ANOVA of groups (frontloaders vs. non-frontloaders vs. water).

#### *Creation of Networks and Identification of Hub Brain Regions*

Key brain regions were identified using a graph theory approach for each group and sex (i.e., 5 networks total were assessed: male frontloaders, male non-frontloaders, male water, female frontloaders, female water). To construct each network, the Pearson correlations between pairs of brain regions, *Figure 2*, were thresholded to only include values above R = 0.7. This allowed the networks and connectivity metrics to be calculated using only strong positive edge weights. The modular structure identified using hierarchical clustering was used to divide each network into modules. The connectivity metrics within module degree z-score (WMDz) and participation coefficient were calculated as described by Guimerà and Nunes Amaral (2005), but modified for networks with weighted edges. WMDz describes how connected a brain region is to the other brain regions within its own module, *Equation 1*. Participation coefficient describes how distributed the connections of a brain region are throughout the network. A participation coefficient of 0 would indicate that a brain region's connections are completely within its own module. A participation coefficient of 1 would indicate that a brain region’s connections are equally distributed among all the modules, *Equation 2*. These connectivity metrics were designed to be applied specifically to modulated networks and can be used to identify each brain region’s contribution to the network, ultimately allowing for the identification of important regions within each module and the network as a whole (55). WMDz and participation coefficient were calculated using customized code from the bctpy Python package ([https://github.com/aestrivex/bctpy](https://github.com/aestrivex/bctpy)), which is a Python version of the widely implemented Matlab brain-connectivity toolbox (56). We thank Dr. Adam Kimbrough’s lab for sharing their customized Python code to calculate these connectivity metrics, see Kimbrough et al. (2020 and 2021) for previous implementation of these analyses. Networks were visualized using Gephi 0.10 and Inkscape 1.1.

Summed weight of all edges between brain region $i$ and the rest of the brain regions in module $s_i$

Average within-module degree of all regions in module $s_i$

$$WMDz = \frac{k_i - \bar{k}_{s_i}}{\sigma_{ks_i}}$$

Standard deviation

*Equation 1. Within Module Degree z-Score.*

Summed weight of all edges between region $i$ and regions in module $s$

$$P_i = 1 - \sum_{s=1}^{N_M} \left( \frac{k_{is}}{k_i} \right)^2$$

Total weight of all edges between brain region $i$ and all other brain regions in the entire network (i.e., total degree)

*Equation 2. Participation Coefficient.*

Once intramodule connectivity (WMDz) and intermodule connectivity (participation coefficient) were identified for each brain region, it was possible to identify the role of each brain region in each network, *Supplementary Figure 3*. The relationship between participation coefficient and WMDz was used to classify brain regions as follows: ultra-peripheral nodes (participation coefficient ≤ 0.05, WMDz < 0.7), peripheral nodes (0.05 < participation coefficient ≤ 0.62, WMDz < 0.7), non-hub connector nodes (0.62 < participation coefficient, WMDz < 0.7), provincial hubs (participation coefficient ≤ 0.3, WMDz > 0.7), and connector hubs (0.3 < participation coefficient, WMDz > 0.7). The participation coefficient divisions are as described in Guimerà and Nunes Amaral (2005), noting that kinless hubs (participation coefficients > 0.8) were not found in our dataset and are not mathematically possible in networks with 3 modules or less (60). The WMDz threshold for hubs versus non-hubs (WMDz > 0.7 classifies a brain region as a hub) is as described in Kimbrough et al. (2020), where there is a need to adjust the threshold for WMDz for c-fos data as compared to functional magnetic resonance imaging data or metabolic network data.

To identify brain regions that may uniquely drive frontloading behavior, lists of connector hubs (high WMDz and high participation coefficient), provincial hubs (high WMDz) and non-hub connector nodes (high participation coefficient) were compared within sex between groups.

Brain regions in these categories with overlap between frontloaders and non-frontloaders or frontloaders and water drinkers were considered to not be uniquely important in frontloading behavior. A final list of connector hubs, provincial hubs, and non-hub connector nodes with no overlap between frontloaders and their within-sex control groups was generated. These brain regions may be uniquely important in frontloading behavior.

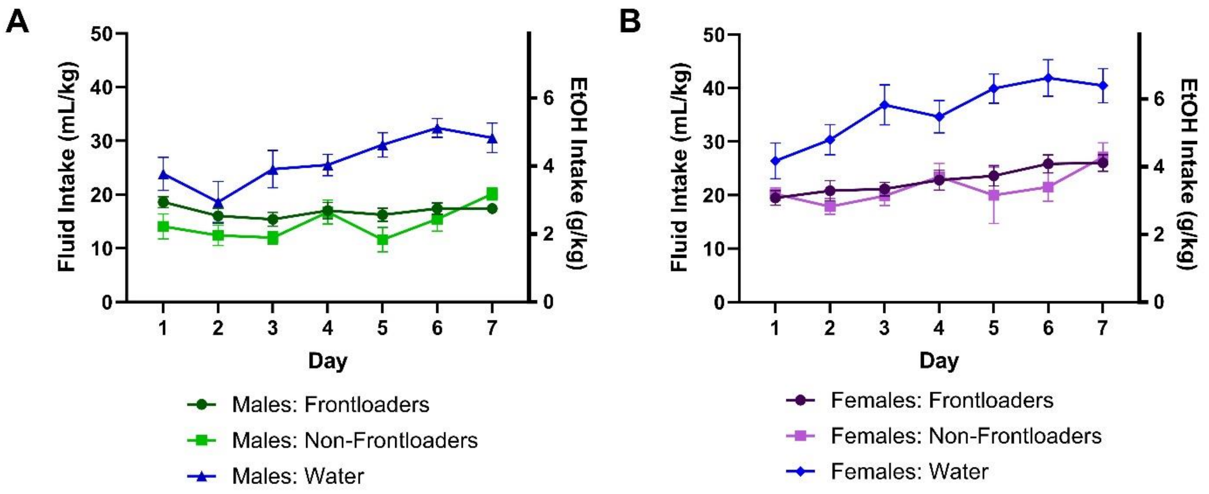


*Supplementary Figure 1.* Total DID session intakes of males (A) and females (B).

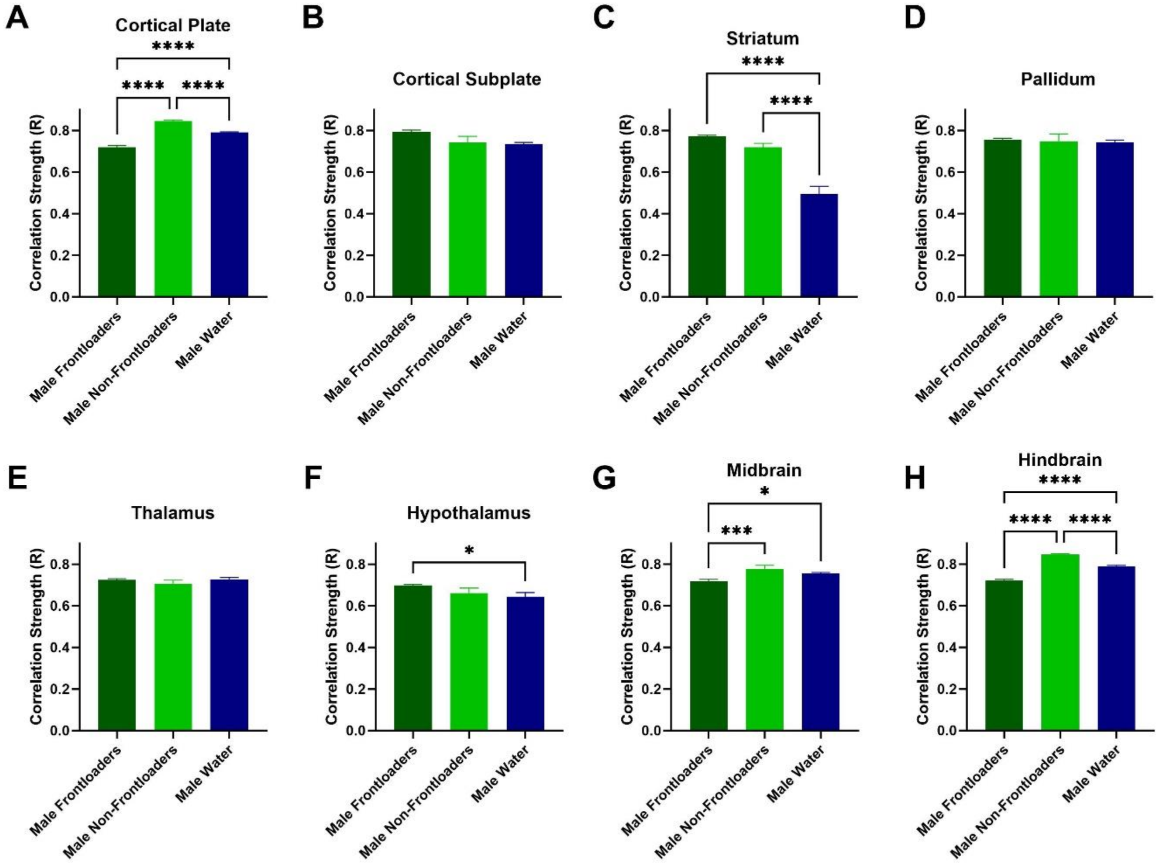


*Supplementary Figure 2.* Correlation strength (R value) is compared across groups within anatomical subdivisions. Note that only significant R values ($p < 0.05$) were included in these analyses. Male frontloaders had lower R values in the cortical plate (A), midbrain (G), and hindbrain (H) as compared to both control groups. Male frontloaders displayed higher R values in the striatum (C) and hypothalamus (F) as compared to water drinkers. These results suggest that frontloading alters the strength of functional connectivity differently across some anatomical subdivisions, and not at all in others.

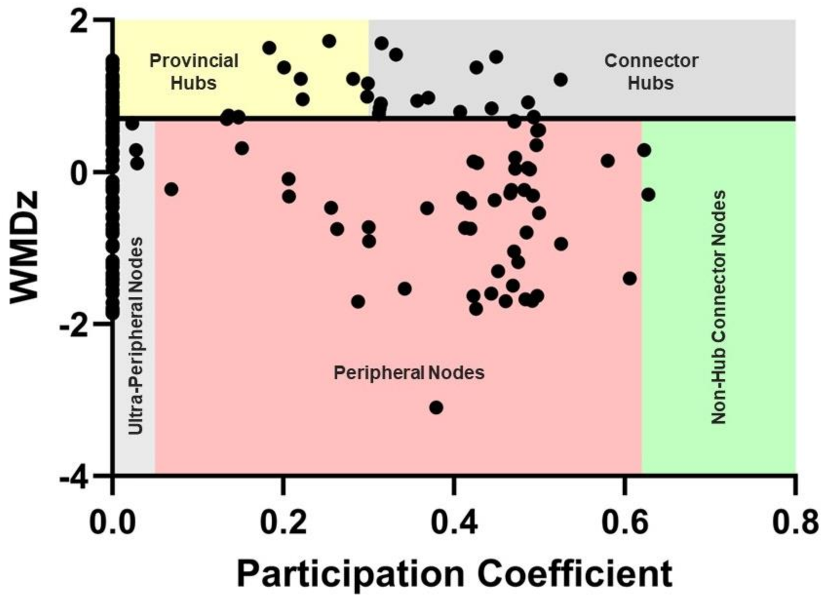


*Supplementary Figure 3.* An example of how the relationship between participation coefficient and within-module degree z-score (WMDz) was used to classify each brain region in a network. Here, male frontloaders are shown. The classifications are as follows: ultra-peripheral nodes (participation coefficient ≤ 0.05, WMDz < 0.7), peripheral nodes (0.05 < participation coefficient ≤ 0.62, WMDz < 0.7), non-hub connector nodes (0.62 < participation coefficient, WMDz < 0.7), provincial hubs (participation coefficient ≤ 0.3, WMDz > 0.7), and connector hubs (0.3 < participation coefficient, WMDz > 0.7).

*Supplementary Table 1.* List of reagents and suppliers for iDISCO tissue clearing and immunohistochemistry.

| Product | Supplier and Product ID |
|---|---|
| Sodium azide | Sigma-Aldrich, S2002-100G |
| TritonX-100 | Fisher, BP151-100 |
| Heparin | Sigma-Aldrich, H3393-50KU |
| Hydrogen Peroxide | Sigma-Aldrich, 216763-100ML |
| Tween-20 | Fisher, BP337-100 |
| Glycine | Sigma-Aldrich, G7126-500G |
| DMSO | Sigma-Aldrich, 472301-1L |
| Methanol | Fisher, BP11054 |
| Dibenzyl Ether | Sigma-Aldrich, 108014-1KG |
| Opaque Black 5 mL Tubes | Dot Scientific, Inc, 229439 |
| C-fos Primary Antibody (Rabbit) | Synaptic Systems, 226 008 |
| Donkey anti-Rabbit IgG (H+L) Highly Cross-Adsorbed Secondary Antibody, Alexa Fluor™ 647 | Thermo-Fisher, # A-31573 |

*Supplementary Table 2.* Intake data considering sex as a factor for experiment 2. Note that the water group is not included in these analyses.

| | | |
|---|---|---|
| **Total 2-Hour DID Session Intake** | | |
| *Source of Variation* | *F (DFn, DFd)* | *P value* |
| Day | F (6, 274) = 4.920 | P < 0.0001**** |
| Sex | F (1, 274) = 76.47 | P < 0.0001**** |
| Group | F (1, 274) = 6.211 | P < 0.05* |
| Day x Sex | F (6, 274) = 0.7349 | P > 0.05 |
| Day x Group | F (6, 274) = 1.221 | P > 0.05 |
| Sex x Group | F (1, 274) = 0.3311 | P > 0.05 |
| Day x Sex x Group | F (6, 274) = 0.4448 | P > 0.05 |
| **Intake in the First 20 Minutes of DID** | | |
| *Source of Variation* | *F (DFn, DFd)* | *P value* |
| Day | F (6, 274) = 3.055 | P < 0.01** |
| Sex | F (1, 274) = 8.399 | P < 0.01** |
| Group | F (1, 274) = 98.43 | P < 0.0001**** |
| Day x Sex | F (6, 274) = 0.2102 | P > 0.05 |
| Day x Group | F (6, 274) = 1.463 | P > 0.05 |
| Sex x Group | F (1, 274) = 2.870 | P > 0.05 |
| Day x Sex x Group | F (6, 274) = 0.3567 | P > 0.05 |
| **Change Point** | | |
| *Source of Variation* | *F (DFn, DFd)* | *P value* |
| Day | F (6, 274) = 0.7275 | P > 0.05 |
| Sex | F (1, 274) = 0.2143 | P > 0.05 |
| Group | F (1, 274) = 442.2 | P < 0.0001**** |
| Day x Sex | F (6, 274) = 2.487 | P < 0.05* |
| Day x Group | F (6, 274) = 0.9241 | P > 0.05 |
| Sex x Group | F (1, 274) = 4.174 | P < 0.05* |
| Day x Sex x Group | F (6, 274) = 2.099 | P > 0.05 |

*Supplementary Table 3.* List of full brain region names and abbreviations.

| Full Brain Region Name | Abbr. | Full Brain Region Name | Abbr. |
|---|---|---|---|
| Frontal pole, cerebral cortex | FRP | Accessory supraoptic group | ASO |
| Somatomotor areas | MO | Paraventricular hypothalamic nucleus | PVH |
| Primary somatosensory area, nose | SSp-n | Anterodorsal preoptic nucleus | ADP |
| Primary somatosensory area, barrel field | SSp-bfd | Anteroventral preoptic nucleus | AVP |
| Primary somatosensory area, lower limb | SSp-ll | Anteroventral periventricular nucleus | AVPV |
| Primary somatosensory area, mouth | SSp-m | Dorsomedial nucleus of the hypothalamus | DMH |
| Primary somatosensory area, upper limb | SSp-ul | Median preoptic nucleus | MEPO |
| Primary somatosensory area, trunk | SSp-tr | Medial preoptic area | MPO |
| Supplemental somatosensory area | SSs | Vascular organ of the lamina terminalis | OV |
| Gustatory areas | GU | Posterodorsal preoptic nucleus | PD |
| Visceral area | VISC | Parastrial nucleus | PS |
| Dorsal auditory area | AUDd | Periventricular hypothalamic nucleus, posterior part | PVp |
| Primary auditory area | AUDp | Periventricular hypothalamic nucleus, preoptic part | PVpo |
| Posterior auditory area | AUDpo | Subparaventricular zone | SBPV |
| Ventral auditory area | AUDv | Suprachiasmatic nucleus | SCH |
| Anterolateral visual area | VISal | Subfornical organ | SFO |
| Anteromedial visual area | VISam | Ventrolateral preoptic nucleus | VLPO |
| Lateral visual area | VISl | Anterior hypothalamic nucleus | AHN |
| Primary visual area | VISp | Mammillary body | MBO |
| Posterolateral visual area | VISpl | Medial preoptic nucleus | MPN |
| posteromedial visual area | VISpm | Paraventricular hypothalamic nucleus, descending division | PVHd |
| Anterior cingulate area, dorsal part | ACAd | Ventromedial hypothalamic nucleus | VMH |
| Anterior cingulate area, ventral part | ACAv | Posterior hypothalamic nucleus | PH |
| Prelimbic area | PL | Lateral hypothalamic area | LHA |
| Infralimbic area | ILA | Lateral preoptic area | LPO |
| Orbital area, lateral part | ORBl | Preparasubthalamic nucleus | PST |
| Orbital area, medial part | ORBm | Parasubthalamic nucleus | PSTN |
| Orbital area, ventrolateral part | ORBvl | Retrochiasmatic area | RCH |
| Agranular insular area, dorsal part | AId | Subthalamic nucleus | STN |
| Agranular insular area, posterior part | AIp | Tuberal nucleus | TU |

Supplementary Table 3 continued

| | | | |
|---|---|---|---|
| Agranular insular area, ventral part | AIv | Zona incerta | ZI |
| Retrosplenial area | RSP | Superior colliculus, sensory related | SCs |
| Temporal association areas | Tea | Inferior colliculus | IC |
| Perirhinal area | PERI | Nucleus of the brachium of the inferior colliculus | NB |
| Ectorhinal area | ECT | Nucleus sagulum | SAG |
| Main olfactory bulb | MOB | Parabigeminal nucleus | PBG |
| Accessory olfactory bulb | AOB | Midbrain trigeminal nucleus | MEV |
| Anterior olfactory nucleus | AON | Substantia nigra, reticular part | SNr |
| Taenia tecta | TT | Ventral tegmental area | VTA |
| Dorsal peduncular area | DP | Midbrain reticular nucleus, retrorubral area | RR |
| Piriform area | PIR | Midbrain reticular nucleus | MRN |
| Cortical amygdalar area | COA | Superior colliculus, motor related | SCm |
| Piriform-amygdalar area | PAA | Periaqueductal gray | PAG |
| Postpiriform transition area | TR | Pretectal region | PRT |
| Field CA1 | CA1 | Cuneiform nucleus | CUN |
| Field CA2 | CA2 | Red nucleus | RN |
| Field CA3 | CA3 | Oculomotor nucleus | III |
| Dentate gyrus | DG | Edinger-Westphal nucleus | EW |
| Entorhinal area | ENT | Trochlear nucleus | IV |
| Parasubiculum | PAR | Ventral tegmental nucleus | VTN |
| Postsubiculum | POST | Anterior tegmental nucleus | AT |
| Presubiculum | PRE | Lateral terminal nucleus of the accessory optic tract | LT |
| Subiculum | SUB | Substantia nigra, compact part | SNc |
| Claustrum | CLA | Pedunculopontine nucleus | PPN |
| Endopiriform nucleus | EP | Midbrain raphe nuclei | Ramb |
| Basolateral amygdalar nucleus | BLA | Nucleus of the lateral lemniscus | NLL |
| Posterior amygdalar nucleus | PA | Principal sensory nucleus of the trigeminal | PSV |
| Caudoputamen | CP | Parabrachial nucleus | PB |
| Nucleus accumbens | ACB | Superior olivary complex | SOC |
| Fundus of striatum | FS | Barringtons nucleus | B |
| Olfactory tubercle | OT | Dorsal tegmental nucleus | DTN |
| Lateral septal nucleus | LS | Pontine central gray | PCG |
| Septofimbrial nucleus | SF | Pontine gray | PG |
| Septohippocampal nucleus | SH | Pontine reticular nucleus, caudal part | PRNc |
| Anterior amygdalar area | AAA | Supragenual nucleus | SG |
| Bed nucleus of the accessory olfactory tract | BA | Supratrigeminal nucleus | SUT |
| Central amygdalar nucleus | CEA | Tegmental reticular nucleus | TRN |
| Intercalated amygdalar nucleus | IA | Motor nucleus of trigeminal | V |

Supplementary Table 3 continued

| | | | |
|---|---|---|---|
| Medial amygdalar nucleus | MEA | Superior central nucleus raphe | CS |
| Pallidum, dorsal region | PALd | Locus ceruleus | LC |
| Pallidum, ventral region | PALv | Laterodorsal tegmental nucleus | LDT |
| Medial septal complex | MSC | Nucleus incertus | NI |
| Pallidum, caudal region | PALc | Pontine reticular nucleus | PRNr |
| Ventral group of the dorsal thalamus | VENT | Nucleus raphe pontis | RPO |
| Ventral posterior complex of the thalamus | VP | Subceruleus nucleus | SLC |
| Subparafascicular nucleus | SPF | Sublaterodorsal nucleus | SLD |
| Subparafascicular area | SPA | Cochlear nuclei | CN |
| Peripeduncular nucleus | PP | Nucleus of the trapezoid body | NTB |
| Geniculate group, dorsal thalamus | GENd | Spinal nucleus of the trigeminal, oral part | SPVO |
| Lateral posterior nucleus of the thalamus | LP | Abducens nucleus | VI |
| Posterior complex of the thalamus | PO | Facial motor nucleus | VII |
| Posterior limiting nucleus of the thalamus | POL | Accessory facial motor nucleus | ACVII |
| Suprageniculate nucleus | SGN | Gigantocellular reticular nucleus | GRN |
| Anteroventral nucleus of thalamus | AV | Intermediate reticular nucleus | IRN |
| Anteromedial nucleus | AM | Magnocellular reticular nucleus | MARN |
| Anterodorsal nucleus | AD | Parvicellular reticular nucleus | PARN |
| Interanteromedial nucleus of the thalamus | IAM | Paragigantocellular reticular nucleus | PGRN |
| Lateral dorsal nucleus of thalamus | LD | Nucleus prepositus | PRP |
| Intermediodorsal nucleus of the thalamus | IMD | Vestibular nuclei | VNC |
| Mediodorsal nucleus of thalamus | MD | Nucleus x | x |
| Submedial nucleus of the thalamus | SMT | Central lobule | CENT |
| Perireunensis nucleus | PR | Culmen | CUL |
| Paraventricular nucleus of the thalamus | PVT | Simple lobule | SIM |
| Parataenial nucleus | PT | Ansiform lobule | AN |
| Nucleus of reuniens | RE | Paramedian lobule | PRM |
| Intralaminar nuclei of the dorsal thalamus | ILM | Paraflocculus | PFL |
| Ventral part of the lateral geniculate complex | LGv | Flocculus | FL |
| Subgeniculate nucleus | SubG | Fastigial nucleus | FN |
| Epithalamus | EPI | Interposed nucleus | IP |
| Supraoptic nucleus | SO | Dentate nucleus | DN |

*Supplementary Table 4.* Each brain region's module and classified role in the male water functional connectivity network.

| Region | Category | Module | PC | WMDz |
|---|---|---|---|---|
| SSp-tr | Connector Hub | 1 | 0.58 | 1.22 |
| SSp-ul | Connector Hub | 1 | 0.58 | 0.94 |
| AIv | Connector Hub | 1 | 0.57 | 1.14 |
| SSp-ll | Connector Hub | 1 | 0.57 | 1.24 |
| LS | Connector Hub | 1 | 0.49 | 1.40 |
| ILA | Connector Hub | 1 | 0.47 | 0.71 |
| ACB | Connector Hub | 1 | 0.43 | 1.20 |
| AId | Peripheral Node | 1 | 0.58 | -0.63 |
| VISam | Peripheral Node | 1 | 0.54 | 0.32 |
| MO | Peripheral Node | 1 | 0.54 | 0.39 |
| GU | Peripheral Node | 1 | 0.50 | -0.66 |
| SH | Peripheral Node | 1 | 0.50 | -1.36 |
| SMT | Peripheral Node | 1 | 0.48 | -1.03 |
| CP | Peripheral Node | 1 | 0.48 | 0.07 |
| SSp-m | Peripheral Node | 1 | 0.45 | -0.65 |
| DP | Peripheral Node | 1 | 0.44 | 0.07 |
| TT | Peripheral Node | 1 | 0.44 | -0.26 |
| VISal | Peripheral Node | 1 | 0.43 | -1.73 |
| ADP | Peripheral Node | 1 | 0.23 | -1.69 |
| FRP | Non-Hub Connector Node | 1 | 0.65 | -0.69 |
| LT | Connector Hub | 2 | 0.62 | 0.71 |
| PBG | Connector Hub | 2 | 0.59 | 1.51 |
| SNr | Connector Hub | 2 | 0.59 | 2.17 |
| MEA | Connector Hub | 2 | 0.58 | 1.72 |
| SSp-n | Connector Hub | 2 | 0.55 | 1.36 |
| SSs | Connector Hub | 2 | 0.54 | 1.48 |
| SO | Provincial Hub | 2 | 0.20 | 1.11 |
| SGN | Peripheral Node | 2 | 0.59 | -0.20 |
| AD | Peripheral Node | 2 | 0.56 | -0.17 |
| AAA | Peripheral Node | 2 | 0.56 | 0.32 |
| VISC | Peripheral Node | 2 | 0.55 | -0.04 |
| LP | Peripheral Node | 2 | 0.54 | 0.60 |
| LPO | Peripheral Node | 2 | 0.53 | 0.57 |
| TU | Peripheral Node | 2 | 0.50 | 0.70 |
| PR | Peripheral Node | 2 | 0.50 | -1.36 |
| AIp | Peripheral Node | 2 | 0.49 | -0.58 |
| VP | Peripheral Node | 2 | 0.48 | 0.26 |
| RSP | Peripheral Node | 2 | 0.45 | -1.01 |
| PALc | Peripheral Node | 2 | 0.45 | 0.54 |

Supplementary Table 4 continued

| | | | | |
|---|---|---|---|---|
| PALv | Peripheral Node | 2 | 0.44 | 0.26 |
| AUDpo | Peripheral Node | 2 | 0.44 | -1.38 |
| BA | Peripheral Node | 2 | 0.43 | 0.32 |
| SF | Peripheral Node | 2 | 0.37 | -0.57 |
| MD | Peripheral Node | 2 | 0.31 | -0.26 |
| PST | Peripheral Node | 2 | 0.30 | -0.18 |
| PS | Ultra-Peripheral Node | 2 | 0.00 | -0.13 |
| SFO | Ultra-Peripheral Node | 2 | 0.00 | -1.34 |
| VISl | Ultra-Peripheral Node | 2 | 0.00 | -1.40 |
| PA | Non-Hub Connector Node | 2 | 0.72 | -1.39 |
| VISpm | Non-Hub Connector Node | 2 | 0.71 | -1.39 |
| VENT | Non-Hub Connector Node | 2 | 0.66 | -0.44 |
| SSp-bfd | Non-Hub Connector Node | 2 | 0.65 | -0.44 |
| CLA | Non-Hub Connector Node | 2 | 0.63 | -1.36 |
| ILM | Connector Hub | 3 | 0.56 | 0.88 |
| MBO | Connector Hub | 3 | 0.53 | 1.22 |
| PALd | Connector Hub | 3 | 0.51 | 1.00 |
| SPF | Connector Hub | 3 | 0.51 | 1.87 |
| MRN | Connector Hub | 3 | 0.44 | 1.23 |
| CA3 | Connector Hub | 3 | 0.43 | 0.88 |
| LD | Connector Hub | 3 | 0.40 | 0.73 |
| AM | Connector Hub | 3 | 0.39 | 1.60 |
| VTA | Connector Hub | 3 | 0.38 | 1.89 |
| STN | Connector Hub | 3 | 0.33 | 1.31 |
| MPN | Provincial Hub | 3 | 0.27 | 0.89 |
| PAA | Provincial Hub | 3 | 0.15 | 1.05 |
| SPA | Peripheral Node | 3 | 0.61 | 0.02 |
| ZI | Peripheral Node | 3 | 0.59 | 0.05 |
| PH | Peripheral Node | 3 | 0.58 | 0.59 |
| PVT | Peripheral Node | 3 | 0.57 | -1.71 |
| FS | Peripheral Node | 3 | 0.57 | -1.42 |
| SOC | Peripheral Node | 3 | 0.57 | -0.66 |
| PVp | Peripheral Node | 3 | 0.57 | 0.43 |
| NB | Peripheral Node | 3 | 0.54 | -0.21 |
| PSTN | Peripheral Node | 3 | 0.53 | 0.24 |
| CEA | Peripheral Node | 3 | 0.52 | 0.67 |
| Tea | Peripheral Node | 3 | 0.50 | -1.43 |
| SubG | Peripheral Node | 3 | 0.50 | -0.79 |
| COA | Peripheral Node | 3 | 0.49 | -1.25 |
| PD | Peripheral Node | 3 | 0.48 | -0.96 |
| AUDd | Peripheral Node | 3 | 0.47 | -0.77 |
| AUDp | Peripheral Node | 3 | 0.45 | -1.98 |

Supplementary Table 4 continued

| | | | | |
|---|---|---|---|---|
| VLPO | Peripheral Node | 3 | 0.45 | -0.39 |
| EP | Peripheral Node | 3 | 0.45 | -0.21 |
| PRT | Peripheral Node | 3 | 0.45 | -0.10 |
| AUDv | Peripheral Node | 3 | 0.44 | -2.12 |
| BLA | Peripheral Node | 3 | 0.43 | 0.19 |
| GENd | Peripheral Node | 3 | 0.43 | -0.22 |
| NTB | Peripheral Node | 3 | 0.42 | -0.68 |
| CA1 | Peripheral Node | 3 | 0.42 | 0.05 |
| PO | Peripheral Node | 3 | 0.41 | 0.25 |
| DG | Peripheral Node | 3 | 0.37 | 0.16 |
| AV | Peripheral Node | 3 | 0.36 | 0.12 |
| DMH | Peripheral Node | 3 | 0.35 | -0.11 |
| PP | Peripheral Node | 3 | 0.23 | -1.25 |
| TR | Peripheral Node | 3 | 0.21 | -1.23 |
| PVHd | Non-Hub Connector Node | 3 | 0.67 | 0.31 |
| CA2 | Non-Hub Connector Node | 3 | 0.65 | -0.16 |
| III | Connector Hub | 4 | 0.41 | 0.77 |
| DTN | Connector Hub | 4 | 0.34 | 1.50 |
| PCG | Connector Hub | 4 | 0.31 | 0.78 |
| TRN | Provincial Hub | 4 | 0.22 | 1.46 |
| NI | Provincial Hub | 4 | 0.21 | 0.96 |
| VISpl | Provincial Hub | 4 | 0.17 | 0.83 |
| FL | Provincial Hub | 4 | 0.11 | 0.97 |
| PRNc | Provincial Hub | 4 | 0.11 | 1.17 |
| PSV | Provincial Hub | 4 | 0.07 | 0.76 |
| LDT | Provincial Hub | 4 | 0.06 | 1.06 |
| V | Provincial Hub | 4 | 0.05 | 1.76 |
| MEV | Provincial Hub | 4 | 0.04 | 2.67 |
| VTN | Peripheral Node | 4 | 0.52 | -0.91 |
| EPI | Peripheral Node | 4 | 0.50 | -2.88 |
| IV | Peripheral Node | 4 | 0.49 | -0.22 |
| SAG | Peripheral Node | 4 | 0.47 | -1.52 |
| IRN | Peripheral Node | 4 | 0.44 | 0.00 |
| PARN | Peripheral Node | 4 | 0.44 | -0.93 |
| RPO | Peripheral Node | 4 | 0.43 | -0.81 |
| CS | Peripheral Node | 4 | 0.39 | 0.60 |
| PGRN | Peripheral Node | 4 | 0.39 | -1.09 |
| VNC | Peripheral Node | 4 | 0.38 | 0.36 |
| AN | Peripheral Node | 4 | 0.37 | 0.28 |
| LC | Peripheral Node | 4 | 0.36 | 0.46 |
| PAG | Peripheral Node | 4 | 0.36 | -0.56 |
| PRP | Peripheral Node | 4 | 0.34 | -1.09 |

Supplementary Table 4 continued

| | | | | |
|---|---|---|---|---|
| SPVO | Peripheral Node | 4 | 0.34 | -1.10 |
| PFL | Peripheral Node | 4 | 0.34 | 0.33 |
| B | Peripheral Node | 4 | 0.33 | -0.51 |
| CN | Peripheral Node | 4 | 0.33 | 0.09 |
| SG | Peripheral Node | 4 | 0.30 | 0.46 |
| GRN | Peripheral Node | 4 | 0.29 | 0.19 |
| PRE | Peripheral Node | 4 | 0.29 | -1.62 |
| VI | Peripheral Node | 4 | 0.27 | 0.37 |
| AT | Peripheral Node | 4 | 0.23 | 0.41 |
| VISp | Peripheral Node | 4 | 0.23 | -1.04 |
| POST | Peripheral Node | 4 | 0.21 | -0.85 |
| SCm | Peripheral Node | 4 | 0.18 | -0.56 |
| RR | Peripheral Node | 4 | 0.15 | -0.04 |
| PG | Peripheral Node | 4 | 0.13 | 0.45 |
| SUT | Peripheral Node | 4 | 0.07 | 0.41 |
| SLD | Ultra-Peripheral Node | 4 | 0.00 | 0.35 |
| SLC | Ultra-Peripheral Node | 4 | 0.00 | -0.24 |
| NLL | Ultra-Peripheral Node | 4 | 0.00 | -0.07 |
| PRNr | Ultra-Peripheral Node | 4 | 0.00 | -0.09 |
| PB | Ultra-Peripheral Node | 4 | 0.00 | -0.13 |
| PPN | Ultra-Peripheral Node | 4 | 0.00 | -0.71 |
| CUN | Ultra-Peripheral Node | 4 | 0.00 | -1.54 |
| SCs | Ultra-Peripheral Node | 4 | 0.00 | -0.90 |
| AOB | Connector Hub | 5 | 0.50 | 0.86 |
| PL | Connector Hub | 5 | 0.49 | 1.93 |
| MOB | Peripheral Node | 5 | 0.56 | -1.11 |
| PIR | Peripheral Node | 5 | 0.55 | 0.42 |
| ACAd | Peripheral Node | 5 | 0.51 | -0.09 |
| ORBm | Peripheral Node | 5 | 0.50 | -0.09 |
| AON | Peripheral Node | 5 | 0.49 | 0.54 |
| ORBvl | Peripheral Node | 5 | 0.49 | 0.08 |
| ORBl | Peripheral Node | 5 | 0.47 | 0.14 |
| ACAv | Ultra-Peripheral Node | 5 | 0.00 | -0.58 |
| ASO | Ultra-Peripheral Node | 5 | 0.00 | -2.11 |
| SIM | Connector Hub | 6 | 0.65 | 1.80 |
| CENT | Connector Hub | 6 | 0.45 | 1.23 |
| ENT | Connector Hub | 6 | 0.54 | 0.81 |
| CUL | Connector Hub | 6 | 0.62 | 1.74 |
| PERI | Peripheral Node | 6 | 0.36 | -0.72 |
| SUB | Peripheral Node | 6 | 0.39 | -0.18 |
| MARN | Peripheral Node | 6 | 0.40 | -0.32 |
| FN | Peripheral Node | 6 | 0.46 | 0.34 |

Supplementary Table 4 continued

| | | | | |
|---|---|---|---|---|
| PRM | Peripheral Node | 6 | 0.46 | 0.42 |
| ACVII | Peripheral Node | 6 | 0.49 | 0.33 |
| IP | Peripheral Node | 6 | 0.49 | 0.51 |
| DN | Peripheral Node | 6 | 0.49 | 0.52 |
| VII | Peripheral Node | 6 | 0.49 | 0.11 |
| EW | Peripheral Node | 6 | 0.50 | -1.28 |
| x | Peripheral Node | 6 | 0.50 | 0.54 |
| PAR | Peripheral Node | 6 | 0.53 | -0.28 |
| IC | Peripheral Node | 6 | 0.58 | 0.23 |
| ECT | Peripheral Node | 6 | 0.60 | -0.75 |
| LHA | Peripheral Node | 6 | 0.66 | -1.32 |
| AHN | Ultra-Peripheral Node | 6 | 0.00 | -2.23 |
| Ramb | Non-Hub Connector Node | 6 | 0.64 | 0.65 |
| LGv | Non-Hub Connector Node | 6 | 0.67 | -1.74 |
| RE | Connector Hub | 7 | 0.50 | 1.19 |
| RCH | Connector Hub | 7 | 0.50 | 0.88 |
| PVpo | Connector Hub | 7 | 0.49 | 0.75 |
| PVH | Connector Hub | 7 | 0.48 | 1.04 |
| MEPO | Connector Hub | 7 | 0.41 | 1.35 |
| AVPV | Connector Hub | 7 | 0.39 | 1.89 |
| IAM | Connector Hub | 7 | 0.35 | 1.02 |
| POL | Peripheral Node | 7 | 0.55 | -0.81 |
| SBPV | Peripheral Node | 7 | 0.54 | -0.11 |
| OT | Peripheral Node | 7 | 0.53 | -0.47 |
| RN | Peripheral Node | 7 | 0.52 | -1.85 |
| SCH | Peripheral Node | 7 | 0.50 | -0.13 |
| IMD | Peripheral Node | 7 | 0.50 | -0.67 |
| OV | Peripheral Node | 7 | 0.48 | -0.21 |
| IA | Peripheral Node | 7 | 0.47 | -1.03 |
| MPO | Peripheral Node | 7 | 0.35 | -1.02 |
| PT | Peripheral Node | 7 | 0.17 | -0.40 |
| AVP | Ultra-Peripheral Node | 7 | 0.00 | -0.15 |
| MSC | Ultra-Peripheral Node | 7 | 0.00 | -1.28 |

*Supplementary Table 5.* Each brain region's module and classified role in the male non-frontloader functional connectivity network.

| Region | Category | Module | PC | WMDz |
|---|---|---|---|---|
| RE | Provincial Hub | 1 | 0.00 | 0.89 |
| PH | Provincial Hub | 1 | 0.00 | 1.18 |
| PVpo | Provincial Hub | 1 | 0.00 | 1.25 |
| PVH | Provincial Hub | 1 | 0.00 | 1.30 |
| AVPV | Provincial Hub | 1 | 0.00 | 1.20 |
| SCH | Provincial Hub | 1 | 0.00 | 0.96 |
| SPA | Provincial Hub | 1 | 0.00 | 0.95 |
| SBPV | Provincial Hub | 1 | 0.00 | 0.75 |
| IMD | Provincial Hub | 1 | 0.00 | 1.45 |
| MEPO | Provincial Hub | 1 | 0.00 | 1.41 |
| TT | Provincial Hub | 1 | 0.00 | 0.77 |
| RPO | Peripheral Node | 1 | 0.50 | -1.43 |
| CS | Peripheral Node | 1 | 0.28 | -1.10 |
| PAG | Peripheral Node | 1 | 0.19 | -0.40 |
| EW | Ultra-Peripheral Node | 1 | 0.00 | -0.77 |
| IV | Ultra-Peripheral Node | 1 | 0.00 | -1.13 |
| MOB | Ultra-Peripheral Node | 1 | 0.00 | -0.08 |
| III | Ultra-Peripheral Node | 1 | 0.00 | -1.34 |
| PVT | Ultra-Peripheral Node | 1 | 0.00 | 0.56 |
| PVp | Ultra-Peripheral Node | 1 | 0.00 | 0.34 |
| AOB | Ultra-Peripheral Node | 1 | 0.00 | -0.28 |
| AON | Ultra-Peripheral Node | 1 | 0.00 | -0.33 |
| OV | Ultra-Peripheral Node | 1 | 0.00 | -0.82 |
| ADP | Ultra-Peripheral Node | 1 | 0.00 | -0.90 |
| AT | Ultra-Peripheral Node | 1 | 0.00 | -1.70 |
| VTN | Ultra-Peripheral Node | 1 | 0.00 | -1.92 |
| SFO | Ultra-Peripheral Node | 1 | 0.00 | 0.12 |
| SH | Ultra-Peripheral Node | 1 | 0.00 | -0.21 |
| Ramb | Ultra-Peripheral Node | 1 | 0.00 | -0.39 |
| FRP | Ultra-Peripheral Node | 1 | 0.00 | -0.34 |
| CUN | Provincial Hub | 2 | 0.06 | 0.88 |
| CA3 | Provincial Hub | 2 | 0.00 | 1.50 |
| CEA | Provincial Hub | 2 | 0.00 | 1.30 |
| ZI | Provincial Hub | 2 | 0.00 | 1.30 |
| PSTN | Provincial Hub | 2 | 0.00 | 1.32 |
| SSp-tr | Provincial Hub | 2 | 0.00 | 1.29 |
| CA2 | Provincial Hub | 2 | 0.00 | 1.26 |
| SSp-bfd | Provincial Hub | 2 | 0.00 | 1.51 |
| PP | Provincial Hub | 2 | 0.00 | 1.44 |

Supplementary Table 5 continued

| | | | | |
|---|---|---|---|---|
| SNc | Provincial Hub | 2 | 0.00 | 1.37 |
| LT | Provincial Hub | 2 | 0.00 | 1.23 |
| PALd | Provincial Hub | 2 | 0.00 | 0.88 |
| CA1 | Provincial Hub | 2 | 0.00 | 1.59 |
| PIR | Provincial Hub | 2 | 0.00 | 1.13 |
| MEA | Provincial Hub | 2 | 0.00 | 0.99 |
| VISC | Provincial Hub | 2 | 0.00 | 0.94 |
| SUB | Provincial Hub | 2 | 0.00 | 1.69 |
| SPF | Provincial Hub | 2 | 0.00 | 1.43 |
| RR | Provincial Hub | 2 | 0.00 | 1.36 |
| TU | Provincial Hub | 2 | 0.00 | 1.31 |
| DG | Provincial Hub | 2 | 0.00 | 1.28 |
| LD | Provincial Hub | 2 | 0.00 | 1.24 |
| SSp-ul | Provincial Hub | 2 | 0.00 | 1.20 |
| AM | Provincial Hub | 2 | 0.00 | 1.14 |
| AAA | Provincial Hub | 2 | 0.00 | 1.06 |
| AIv | Provincial Hub | 2 | 0.00 | 1.01 |
| AIp | Provincial Hub | 2 | 0.00 | 0.98 |
| OT | Provincial Hub | 2 | 0.00 | 0.92 |
| LPO | Provincial Hub | 2 | 0.00 | 0.89 |
| MD | Provincial Hub | 2 | 0.00 | 0.78 |
| IA | Provincial Hub | 2 | 0.00 | 0.75 |
| ASO | Provincial Hub | 2 | 0.00 | 0.71 |
| MO | Provincial Hub | 2 | 0.00 | 1.18 |
| LP | Provincial Hub | 2 | 0.00 | 1.14 |
| PO | Provincial Hub | 2 | 0.00 | 1.50 |
| VENT | Provincial Hub | 2 | 0.00 | 1.09 |
| VTA | Provincial Hub | 2 | 0.00 | 1.04 |
| SSp-ll | Provincial Hub | 2 | 0.00 | 0.99 |
| ORBl | Provincial Hub | 2 | 0.00 | 0.94 |
| CLA | Provincial Hub | 2 | 0.00 | 0.90 |
| LHA | Provincial Hub | 2 | 0.00 | 0.86 |
| PALv | Provincial Hub | 2 | 0.00 | 0.71 |
| ECT | Provincial Hub | 2 | 0.00 | 1.06 |
| BLA | Provincial Hub | 2 | 0.00 | 0.98 |
| VP | Provincial Hub | 2 | 0.00 | 0.93 |
| STN | Provincial Hub | 2 | 0.00 | 0.87 |
| RSP | Provincial Hub | 2 | 0.00 | 1.57 |
| SNr | Provincial Hub | 2 | 0.00 | 1.45 |
| PERI | Provincial Hub | 2 | 0.00 | 0.81 |
| MRN | Provincial Hub | 2 | 0.00 | 0.74 |
| MBO | Peripheral Node | 2 | 0.44 | -1.98 |

Supplementary Table 5 continued

| | | | | |
|---|---|---|---|---|
| PRP | Peripheral Node | 2 | 0.41 | -1.44 |
| PGRN | Peripheral Node | 2 | 0.40 | -1.33 |
| IRN | Peripheral Node | 2 | 0.36 | -1.21 |
| CN | Peripheral Node | 2 | 0.35 | -1.36 |
| NTB | Peripheral Node | 2 | 0.33 | -1.78 |
| PARN | Peripheral Node | 2 | 0.31 | -1.09 |
| GRN | Peripheral Node | 2 | 0.30 | -0.94 |
| SAG | Peripheral Node | 2 | 0.28 | -1.88 |
| DTN | Peripheral Node | 2 | 0.26 | -1.65 |
| DN | Peripheral Node | 2 | 0.24 | -1.13 |
| MSC | Peripheral Node | 2 | 0.21 | -1.82 |
| V | Peripheral Node | 2 | 0.21 | -0.76 |
| MARN | Peripheral Node | 2 | 0.17 | -1.05 |
| VI | Peripheral Node | 2 | 0.16 | -0.53 |
| VNC | Peripheral Node | 2 | 0.12 | -0.01 |
| AN | Peripheral Node | 2 | 0.11 | -0.49 |
| VISpl | Peripheral Node | 2 | 0.11 | -1.02 |
| VISl | Peripheral Node | 2 | 0.11 | -1.54 |
| LC | Peripheral Node | 2 | 0.09 | -0.02 |
| VISp | Peripheral Node | 2 | 0.09 | -0.20 |
| SG | Peripheral Node | 2 | 0.08 | 0.16 |
| VII | Peripheral Node | 2 | 0.08 | -1.37 |
| PFL | Peripheral Node | 2 | 0.06 | -0.22 |
| SOC | Peripheral Node | 2 | 0.06 | -1.10 |
| FL | Peripheral Node | 2 | 0.06 | 0.08 |
| ENT | Ultra-Peripheral Node | 2 | 0.05 | -0.69 |
| BA | Ultra-Peripheral Node | 2 | 0.00 | 0.63 |
| PALc | Ultra-Peripheral Node | 2 | 0.00 | 0.47 |
| AUDp | Ultra-Peripheral Node | 2 | 0.00 | -0.51 |
| EP | Ultra-Peripheral Node | 2 | 0.00 | 0.47 |
| POST | Ultra-Peripheral Node | 2 | 0.00 | 0.35 |
| SF | Ultra-Peripheral Node | 2 | 0.00 | 0.21 |
| DMH | Ultra-Peripheral Node | 2 | 0.00 | -0.97 |
| CP | Ultra-Peripheral Node | 2 | 0.00 | 0.54 |
| VISpm | Ultra-Peripheral Node | 2 | 0.00 | -0.10 |
| PB | Ultra-Peripheral Node | 2 | 0.00 | -0.31 |
| AUDd | Ultra-Peripheral Node | 2 | 0.00 | -0.10 |
| VISal | Ultra-Peripheral Node | 2 | 0.00 | -0.75 |
| PPN | Ultra-Peripheral Node | 2 | 0.00 | -0.99 |
| Tea | Ultra-Peripheral Node | 2 | 0.00 | 0.55 |
| ILM | Ultra-Peripheral Node | 2 | 0.00 | 0.42 |
| GENd | Ultra-Peripheral Node | 2 | 0.00 | 0.02 |

Supplementary Table 5 continued

| | | | | |
|---|---|---|---|---|
| PL | Ultra-Peripheral Node | 2 | 0.00 | -0.08 |
| CENT | Ultra-Peripheral Node | 2 | 0.00 | -0.09 |
| PCG | Ultra-Peripheral Node | 2 | 0.00 | -0.43 |
| PS | Ultra-Peripheral Node | 2 | 0.00 | -0.66 |
| PR | Ultra-Peripheral Node | 2 | 0.00 | -0.71 |
| LDT | Ultra-Peripheral Node | 2 | 0.00 | -1.04 |
| EPI | Ultra-Peripheral Node | 2 | 0.00 | -1.16 |
| SLD | Ultra-Peripheral Node | 2 | 0.00 | -1.35 |
| ILA | Ultra-Peripheral Node | 2 | 0.00 | 0.30 |
| LS | Ultra-Peripheral Node | 2 | 0.00 | 0.30 |
| PST | Ultra-Peripheral Node | 2 | 0.00 | -0.19 |
| AUDv | Ultra-Peripheral Node | 2 | 0.00 | -0.56 |
| SSp-m | Ultra-Peripheral Node | 2 | 0.00 | -0.89 |
| PT | Ultra-Peripheral Node | 2 | 0.00 | -1.23 |
| FS | Ultra-Peripheral Node | 2 | 0.00 | 0.69 |
| ACB | Ultra-Peripheral Node | 2 | 0.00 | 0.66 |
| AV | Ultra-Peripheral Node | 2 | 0.00 | 0.58 |
| SGN | Ultra-Peripheral Node | 2 | 0.00 | 0.10 |
| SSp-n | Ultra-Peripheral Node | 2 | 0.00 | -0.14 |
| ACAd | Ultra-Peripheral Node | 2 | 0.00 | -0.25 |
| MPO | Ultra-Peripheral Node | 2 | 0.00 | -0.70 |
| NLL | Ultra-Peripheral Node | 2 | 0.00 | -0.71 |
| LGv | Ultra-Peripheral Node | 2 | 0.00 | -1.36 |
| AId | Ultra-Peripheral Node | 2 | 0.00 | 0.60 |
| PD | Ultra-Peripheral Node | 2 | 0.00 | 0.57 |
| SSs | Ultra-Peripheral Node | 2 | 0.00 | 0.43 |
| DP | Ultra-Peripheral Node | 2 | 0.00 | 0.24 |
| SCm | Ultra-Peripheral Node | 2 | 0.00 | 0.09 |
| COA | Ultra-Peripheral Node | 2 | 0.00 | 0.09 |
| RCH | Ultra-Peripheral Node | 2 | 0.00 | -0.03 |
| POL | Ultra-Peripheral Node | 2 | 0.00 | -0.18 |
| B | Ultra-Peripheral Node | 2 | 0.00 | -0.42 |
| IC | Ultra-Peripheral Node | 2 | 0.00 | -0.47 |
| PBG | Ultra-Peripheral Node | 2 | 0.00 | -0.62 |
| TR | Ultra-Peripheral Node | 2 | 0.00 | -0.96 |
| PRT | Ultra-Peripheral Node | 2 | 0.00 | -1.02 |
| ACAv | Ultra-Peripheral Node | 2 | 0.00 | -1.10 |
| PVHd | Ultra-Peripheral Node | 2 | 0.00 | -1.31 |
| SubG | Ultra-Peripheral Node | 2 | 0.00 | -1.63 |
| IAM | Ultra-Peripheral Node | 2 | 0.00 | -1.69 |
| PRNr | Ultra-Peripheral Node | 2 | 0.00 | -1.77 |
| VMH | Ultra-Peripheral Node | 2 | 0.00 | -1.78 |

Supplementary Table 5 continued

| | | | | |
|---|---|---|---|---|
| AVP | Ultra-Peripheral Node | 2 | 0.00 | -1.91 |
| ORBm | Ultra-Peripheral Node | 2 | 0.00 | -2.01 |
| NB | Ultra-Peripheral Node | 2 | 0.00 | 0.53 |
| SCs | Ultra-Peripheral Node | 2 | 0.00 | 0.01 |
| SIM | Ultra-Peripheral Node | 2 | 0.00 | -0.73 |
| SMT | Ultra-Peripheral Node | 2 | 0.00 | -1.18 |
| SLC | Ultra-Peripheral Node | 2 | 0.00 | -1.39 |
| PRE | Ultra-Peripheral Node | 2 | 0.00 | 0.69 |
| GU | Ultra-Peripheral Node | 2 | 0.00 | 0.56 |
| MEV | Ultra-Peripheral Node | 2 | 0.00 | 0.07 |
| RN | Ultra-Peripheral Node | 2 | 0.00 | -0.11 |
| PAR | Ultra-Peripheral Node | 2 | 0.00 | -0.49 |
| CUL | Ultra-Peripheral Node | 2 | 0.00 | -0.49 |
| SO | Ultra-Peripheral Node | 2 | 0.00 | -0.59 |
| AD | Ultra-Peripheral Node | 2 | 0.00 | -1.15 |
| MPN | Ultra-Peripheral Node | 2 | 0.00 | -1.91 |
| VLPO | Ultra-Peripheral Node | 2 | 0.00 | 0.53 |
| PA | Ultra-Peripheral Node | 2 | 0.00 | 0.31 |
| AUDpo | Ultra-Peripheral Node | 2 | 0.00 | -0.29 |
| NI | Ultra-Peripheral Node | 2 | 0.00 | -0.76 |
| PAA | Ultra-Peripheral Node | 2 | 0.00 | 0.58 |
| AHN | Ultra-Peripheral Node | 2 | 0.00 | 0.13 |
| ORBvl | Ultra-Peripheral Node | 2 | 0.00 | -0.59 |
| VISam | Ultra-Peripheral Node | 2 | 0.00 | 0.11 |
| IP | Connector Hub | 3 | 0.43 | 1.21 |
| SPVO | Connector Hub | 3 | 0.42 | 1.21 |
| PRM | Connector Hub | 3 | 0.40 | 1.10 |
| PSV | Peripheral Node | 3 | 0.52 | -0.07 |
| SUT | Peripheral Node | 3 | 0.52 | -0.03 |
| x | Peripheral Node | 3 | 0.49 | -0.23 |
| FN | Peripheral Node | 3 | 0.47 | -0.22 |
| TRN | Peripheral Node | 3 | 0.43 | -1.42 |
| PG | Peripheral Node | 3 | 0.41 | 0.35 |
| PRNc | Peripheral Node | 3 | 0.34 | -1.91 |

*Supplementary Table 6.* Each brain region's module and classified role in the male frontloader functional connectivity network.

| Region | Category | Module | PC | WMDz |
|---|---|---|---|---|
| CP | Provincial Hub | 1 | 0.00 | 1.42 |
| AIv | Provincial Hub | 1 | 0.00 | 1.23 |
| CA3 | Provincial Hub | 1 | 0.00 | 1.23 |
| STN | Provincial Hub | 1 | 0.00 | 1.37 |
| PAA | Provincial Hub | 1 | 0.00 | 1.04 |
| RSP | Provincial Hub | 1 | 0.00 | 0.84 |
| RN | Provincial Hub | 1 | 0.00 | 0.78 |
| BLA | Provincial Hub | 1 | 0.00 | 0.78 |
| AAA | Provincial Hub | 1 | 0.00 | 0.77 |
| LS | Provincial Hub | 1 | 0.00 | 0.75 |
| SSp-tr | Provincial Hub | 1 | 0.00 | 1.21 |
| EP | Provincial Hub | 1 | 0.00 | 1.16 |
| CA1 | Provincial Hub | 1 | 0.00 | 1.14 |
| PALc | Provincial Hub | 1 | 0.00 | 0.83 |
| VENT | Provincial Hub | 1 | 0.00 | 0.81 |
| SSp-ul | Provincial Hub | 1 | 0.00 | 1.47 |
| MO | Provincial Hub | 1 | 0.00 | 1.35 |
| SSp-ll | Provincial Hub | 1 | 0.00 | 1.26 |
| ACB | Provincial Hub | 1 | 0.00 | 1.12 |
| SCm | Provincial Hub | 1 | 0.00 | 1.02 |
| PIR | Provincial Hub | 1 | 0.00 | 0.99 |
| LPO | Provincial Hub | 1 | 0.00 | 0.98 |
| SSp-bfd | Provincial Hub | 1 | 0.00 | 0.97 |
| VISC | Provincial Hub | 1 | 0.00 | 0.92 |
| LHA | Provincial Hub | 1 | 0.00 | 0.86 |
| MEA | Provincial Hub | 1 | 0.00 | 0.76 |
| SNr | Provincial Hub | 1 | 0.00 | 0.75 |
| SNc | Provincial Hub | 1 | 0.00 | 1.33 |
| AON | Provincial Hub | 1 | 0.00 | 0.84 |
| MRN | Provincial Hub | 1 | 0.00 | 1.09 |
| PALd | Provincial Hub | 1 | 0.00 | 1.05 |
| AIp | Provincial Hub | 1 | 0.00 | 0.99 |
| COA | Provincial Hub | 1 | 0.00 | 0.92 |
| PRE | Provincial Hub | 1 | 0.00 | 0.85 |
| CEA | Provincial Hub | 1 | 0.00 | 0.84 |
| PALv | Provincial Hub | 1 | 0.00 | 1.23 |
| AId | Provincial Hub | 1 | 0.00 | 1.01 |
| CN | Peripheral Node | 1 | 0.50 | -1.63 |
| PD | Peripheral Node | 1 | 0.49 | -1.69 |

Supplementary Table 6 continued

| | | | | |
|---|---|---|---|---|
| VTA | Peripheral Node | 1 | 0.48 | -1.67 |
| TT | Peripheral Node | 1 | 0.47 | -1.49 |
| RCH | Peripheral Node | 1 | 0.46 | -1.70 |
| FN | Peripheral Node | 1 | 0.44 | -1.60 |
| V | Peripheral Node | 1 | 0.43 | -1.80 |
| AHN | Peripheral Node | 1 | 0.42 | -1.63 |
| PSV | Peripheral Node | 1 | 0.34 | -1.53 |
| PRNr | Peripheral Node | 1 | 0.29 | -1.70 |
| MSC | Peripheral Node | 1 | 0.07 | -0.22 |
| ENT | Ultra-Peripheral Node | 1 | 0.03 | 0.12 |
| SPF | Ultra-Peripheral Node | 1 | 0.03 | 0.29 |
| SUB | Ultra-Peripheral Node | 1 | 0.02 | 0.64 |
| DP | Ultra-Peripheral Node | 1 | 0.00 | 0.55 |
| POL | Ultra-Peripheral Node | 1 | 0.00 | 0.57 |
| AM | Ultra-Peripheral Node | 1 | 0.00 | 0.41 |
| PO | Ultra-Peripheral Node | 1 | 0.00 | 0.38 |
| DG | Ultra-Peripheral Node | 1 | 0.00 | -0.24 |
| VP | Ultra-Peripheral Node | 1 | 0.00 | 0.57 |
| AUDpo | Ultra-Peripheral Node | 1 | 0.00 | -0.17 |
| AUDd | Ultra-Peripheral Node | 1 | 0.00 | 0.65 |
| CA2 | Ultra-Peripheral Node | 1 | 0.00 | 0.54 |
| BA | Ultra-Peripheral Node | 1 | 0.00 | 0.54 |
| ORBvl | Ultra-Peripheral Node | 1 | 0.00 | -0.02 |
| SSp-m | Ultra-Peripheral Node | 1 | 0.00 | 0.64 |
| AD | Ultra-Peripheral Node | 1 | 0.00 | -1.17 |
| CUN | Ultra-Peripheral Node | 1 | 0.00 | -1.35 |
| PA | Ultra-Peripheral Node | 1 | 0.00 | 0.45 |
| VISpl | Ultra-Peripheral Node | 1 | 0.00 | -0.79 |
| AOB | Ultra-Peripheral Node | 1 | 0.00 | -0.95 |
| ACAd | Ultra-Peripheral Node | 1 | 0.00 | -0.60 |
| SubG | Ultra-Peripheral Node | 1 | 0.00 | -0.98 |
| SAG | Ultra-Peripheral Node | 1 | 0.00 | -1.33 |
| PAR | Ultra-Peripheral Node | 1 | 0.00 | -1.55 |
| OT | Ultra-Peripheral Node | 1 | 0.00 | 0.60 |
| PERI | Ultra-Peripheral Node | 1 | 0.00 | 0.48 |
| GU | Ultra-Peripheral Node | 1 | 0.00 | 0.66 |
| PP | Ultra-Peripheral Node | 1 | 0.00 | 0.66 |
| ILM | Ultra-Peripheral Node | 1 | 0.00 | 0.65 |
| LD | Ultra-Peripheral Node | 1 | 0.00 | 0.42 |
| POST | Ultra-Peripheral Node | 1 | 0.00 | 0.24 |
| Tea | Ultra-Peripheral Node | 1 | 0.00 | 0.07 |
| AUDv | Ultra-Peripheral Node | 1 | 0.00 | -0.12 |

Supplementary Table 6 continued

| | | | | |
|---|---|---|---|---|
| PSTN | Ultra-Peripheral Node | 1 | 0.00 | -0.15 |
| SF | Ultra-Peripheral Node | 1 | 0.00 | -0.20 |
| AUDp | Ultra-Peripheral Node | 1 | 0.00 | -0.35 |
| VISam | Ultra-Peripheral Node | 1 | 0.00 | -0.40 |
| ORBm | Ultra-Peripheral Node | 1 | 0.00 | -0.46 |
| LP | Ultra-Peripheral Node | 1 | 0.00 | -0.57 |
| ILA | Ultra-Peripheral Node | 1 | 0.00 | -0.62 |
| VISl | Ultra-Peripheral Node | 1 | 0.00 | -0.70 |
| SGN | Ultra-Peripheral Node | 1 | 0.00 | -0.75 |
| TU | Ultra-Peripheral Node | 1 | 0.00 | -0.81 |
| NB | Ultra-Peripheral Node | 1 | 0.00 | -1.17 |
| ACAv | Ultra-Peripheral Node | 1 | 0.00 | -1.25 |
| PPN | Ultra-Peripheral Node | 1 | 0.00 | -1.40 |
| PBG | Ultra-Peripheral Node | 1 | 0.00 | -1.42 |
| LGv | Ultra-Peripheral Node | 1 | 0.00 | -1.47 |
| PRT | Ultra-Peripheral Node | 1 | 0.00 | -1.51 |
| NLL | Ultra-Peripheral Node | 1 | 0.00 | -1.72 |
| SCs | Ultra-Peripheral Node | 1 | 0.00 | -1.77 |
| ASO | Ultra-Peripheral Node | 1 | 0.00 | -1.79 |
| PST | Ultra-Peripheral Node | 1 | 0.00 | -1.83 |
| SPVO | Ultra-Peripheral Node | 1 | 0.00 | -1.86 |
| SSp-n | Ultra-Peripheral Node | 1 | 0.00 | 0.65 |
| SSs | Ultra-Peripheral Node | 1 | 0.00 | 0.63 |
| FS | Ultra-Peripheral Node | 1 | 0.00 | 0.60 |
| ZI | Ultra-Peripheral Node | 1 | 0.00 | 0.56 |
| RR | Ultra-Peripheral Node | 1 | 0.00 | 0.51 |
| VISpm | Ultra-Peripheral Node | 1 | 0.00 | 0.35 |
| ECT | Ultra-Peripheral Node | 1 | 0.00 | 0.20 |
| PL | Ultra-Peripheral Node | 1 | 0.00 | 0.13 |
| AV | Ultra-Peripheral Node | 1 | 0.00 | 0.00 |
| SO | Ultra-Peripheral Node | 1 | 0.00 | -1.28 |
| CLA | Ultra-Peripheral Node | 1 | 0.00 | 0.58 |
| VISal | Ultra-Peripheral Node | 1 | 0.00 | 0.19 |
| GENd | Ultra-Peripheral Node | 1 | 0.00 | 0.07 |
| VISp | Ultra-Peripheral Node | 1 | 0.00 | -0.26 |
| MOB | Ultra-Peripheral Node | 1 | 0.00 | -0.32 |
| FRP | Ultra-Peripheral Node | 1 | 0.00 | -1.08 |
| LT | Ultra-Peripheral Node | 1 | 0.00 | -1.51 |
| IA | Ultra-Peripheral Node | 1 | 0.00 | 0.56 |
| ORBl | Ultra-Peripheral Node | 1 | 0.00 | 0.10 |
| MD | Ultra-Peripheral Node | 1 | 0.00 | -1.00 |
| TR | Ultra-Peripheral Node | 1 | 0.00 | 0.31 |

Supplementary Table 6 continued

| | | | | |
|---|---|---|---|---|
| PG | Connector Hub | 2 | 0.53 | 1.21 |
| SBPV | Connector Hub | 2 | 0.49 | 0.72 |
| RPO | Connector Hub | 2 | 0.49 | 0.91 |
| SG | Connector Hub | 2 | 0.41 | 0.79 |
| MEPO | Connector Hub | 2 | 0.37 | 0.97 |
| PVHd | Connector Hub | 2 | 0.36 | 0.93 |
| MARN | Connector Hub | 2 | 0.33 | 1.54 |
| AVPV | Connector Hub | 2 | 0.32 | 1.69 |
| PVpo | Connector Hub | 2 | 0.31 | 0.90 |
| CUL | Connector Hub | 2 | 0.31 | 0.83 |
| SIM | Connector Hub | 2 | 0.31 | 0.76 |
| GRN | Provincial Hub | 2 | 0.30 | 1.16 |
| PFL | Provincial Hub | 2 | 0.30 | 0.99 |
| FL | Provincial Hub | 2 | 0.28 | 1.22 |
| VNC | Provincial Hub | 2 | 0.25 | 1.72 |
| MEV | Provincial Hub | 2 | 0.22 | 0.95 |
| SCH | Provincial Hub | 2 | 0.22 | 1.22 |
| LC | Provincial Hub | 2 | 0.20 | 1.37 |
| PARN | Provincial Hub | 2 | 0.15 | 0.72 |
| IRN | Provincial Hub | 2 | 0.14 | 0.74 |
| VII | Provincial Hub | 2 | 0.13 | 0.70 |
| MPO | Non-hub Conn. Node | 2 | 0.63 | -0.29 |
| PS | Non-hub Conn. Node | 2 | 0.62 | 0.29 |
| SH | Peripheral Node | 2 | 0.61 | -1.40 |
| MBO | Peripheral Node | 2 | 0.58 | 0.15 |
| AT | Peripheral Node | 2 | 0.53 | -0.94 |
| VI | Peripheral Node | 2 | 0.50 | -0.54 |
| EW | Peripheral Node | 2 | 0.50 | 0.55 |
| IMD | Peripheral Node | 2 | 0.50 | 0.36 |
| PRP | Peripheral Node | 2 | 0.49 | -0.31 |
| LDT | Peripheral Node | 2 | 0.48 | -1.18 |
| PAG | Peripheral Node | 2 | 0.47 | -1.04 |
| VMH | Peripheral Node | 2 | 0.42 | -0.74 |
| PB | Peripheral Node | 2 | 0.41 | -0.73 |
| NTB | Peripheral Node | 2 | 0.41 | -0.34 |
| SLD | Peripheral Node | 2 | 0.30 | -0.91 |
| SUT | Peripheral Node | 2 | 0.30 | -0.72 |
| DN | Peripheral Node | 2 | 0.26 | -0.75 |
| B | Peripheral Node | 2 | 0.26 | -0.47 |
| IP | Peripheral Node | 2 | 0.21 | -0.32 |
| PCG | Peripheral Node | 2 | 0.21 | -0.09 |
| OV | Peripheral Node | 2 | 0.15 | 0.32 |

Supplementary Table 6 continued

| | | | | |
|---|---|---|---|---|
| CENT | Ultra-Peripheral Node | 2 | 0.00 | 0.16 |
| AN | Ultra-Peripheral Node | 2 | 0.00 | 0.43 |
| SOC | Ultra-Peripheral Node | 2 | 0.00 | 0.27 |
| SLC | Ultra-Peripheral Node | 2 | 0.00 | -0.98 |
| PGRN | Ultra-Peripheral Node | 2 | 0.00 | -1.20 |
| PRM | Ultra-Peripheral Node | 2 | 0.00 | -1.33 |
| x | Ultra-Peripheral Node | 2 | 0.00 | -1.33 |
| IC | Ultra-Peripheral Node | 2 | 0.00 | -1.40 |
| EPI | Ultra-Peripheral Node | 2 | 0.00 | -1.40 |
| SFO | Ultra-Peripheral Node | 2 | 0.00 | -1.40 |
| SMT | Ultra-Peripheral Node | 2 | 0.00 | -1.40 |
| PR | Ultra-Peripheral Node | 2 | 0.00 | -1.60 |
| IAM | Ultra-Peripheral Node | 2 | 0.00 | -1.81 |
| PVH | Connector Hub | 3 | 0.45 | 1.51 |
| RE | Connector Hub | 3 | 0.44 | 0.83 |
| TRN | Connector Hub | 3 | 0.43 | 1.37 |
| PVT | Provincial Hub | 3 | 0.18 | 1.63 |
| VTN | Peripheral Node | 3 | 0.50 | 0.56 |
| PVp | Peripheral Node | 3 | 0.49 | 0.04 |
| CS | Peripheral Node | 3 | 0.49 | 0.06 |
| DTN | Peripheral Node | 3 | 0.48 | -0.79 |
| SPA | Peripheral Node | 3 | 0.48 | -0.23 |
| MPN | Peripheral Node | 3 | 0.47 | 0.19 |
| ADP | Peripheral Node | 3 | 0.47 | 0.05 |
| Ramb | Peripheral Node | 3 | 0.47 | 0.67 |
| III | Peripheral Node | 3 | 0.47 | -0.23 |
| IV | Peripheral Node | 3 | 0.47 | -0.28 |
| NI | Peripheral Node | 3 | 0.45 | -1.30 |
| AVP | Peripheral Node | 3 | 0.45 | -0.37 |
| DMH | Peripheral Node | 3 | 0.43 | 0.12 |
| PH | Peripheral Node | 3 | 0.42 | 0.14 |
| PT | Peripheral Node | 3 | 0.42 | -0.41 |
| PRNc | Peripheral Node | 3 | 0.38 | -3.10 |
| VLPO | Peripheral Node | 3 | 0.37 | -0.47 |

*Supplementary Table 7.* Each brain region's module and classified role in the female water drinking functional connectivity network.

| Region | Category | Module | PC | WMDz |
|---|---|---|---|---|
| PRP | Connector Hub | 1 | 0.36 | 0.89 |
| ACVII | Connector Hub | 1 | 0.36 | 0.86 |
| MEPO | Connector Hub | 1 | 0.45 | 1.44 |
| PGRN | Connector Hub | 1 | 0.46 | 1.15 |
| SBPV | Connector Hub | 1 | 0.48 | 1.73 |
| GRN | Connector Hub | 1 | 0.50 | 1.09 |
| SG | Connector Hub | 1 | 0.50 | 1.06 |
| VI | Connector Hub | 1 | 0.50 | 1.05 |
| NTB | Connector Hub | 1 | 0.54 | 2.46 |
| SFO | Connector Hub | 1 | 0.56 | 0.85 |
| MARN | Connector Hub | 1 | 0.56 | 2.02 |
| GU | Connector Hub | 1 | 0.59 | 1.12 |
| SCH | Provincial Hub | 1 | 0.00 | 1.08 |
| AVPV | Provincial Hub | 1 | 0.13 | 1.83 |
| PB | Peripheral Node | 1 | 0.22 | -1.36 |
| OV | Peripheral Node | 1 | 0.29 | -0.30 |
| PVpo | Peripheral Node | 1 | 0.29 | -0.30 |
| VISam | Peripheral Node | 1 | 0.30 | -1.33 |
| SSs | Peripheral Node | 1 | 0.38 | -0.19 |
| PVp | Peripheral Node | 1 | 0.44 | 0.23 |
| PD | Peripheral Node | 1 | 0.45 | -1.09 |
| IMD | Peripheral Node | 1 | 0.46 | -0.37 |
| EW | Peripheral Node | 1 | 0.46 | -0.39 |
| SH | Peripheral Node | 1 | 0.47 | 0.18 |
| OT | Peripheral Node | 1 | 0.47 | -0.45 |
| SPF | Peripheral Node | 1 | 0.47 | -0.91 |
| SSp-n | Peripheral Node | 1 | 0.47 | 0.29 |
| PSTN | Peripheral Node | 1 | 0.47 | -0.24 |
| SF | Peripheral Node | 1 | 0.48 | -0.93 |
| RPO | Peripheral Node | 1 | 0.48 | -0.35 |
| TU | Peripheral Node | 1 | 0.48 | 0.42 |
| SNc | Peripheral Node | 1 | 0.49 | -0.71 |
| SSp-tr | Peripheral Node | 1 | 0.49 | -0.02 |
| SSp-m | Peripheral Node | 1 | 0.49 | 0.54 |
| FRP | Peripheral Node | 1 | 0.49 | -0.83 |
| AV | Peripheral Node | 1 | 0.52 | -0.08 |
| VISpm | Peripheral Node | 1 | 0.53 | -0.22 |
| FS | Peripheral Node | 1 | 0.55 | -0.25 |
| AId | Peripheral Node | 1 | 0.55 | 0.03 |

Supplementary Table 7 continued

| | | | | |
|---|---|---|---|---|
| CLA | Peripheral Node | 1 | 0.57 | -1.10 |
| SUT | Peripheral Node | 1 | 0.59 | -0.25 |
| CEA | Peripheral Node | 1 | 0.59 | -0.19 |
| GENd | Ultra-Peripheral Node | 1 | 0.00 | -1.57 |
| SSp-bfd | Ultra-Peripheral Node | 1 | 0.00 | -1.15 |
| MEV | Ultra-Peripheral Node | 1 | 0.00 | -1.11 |
| RR | Ultra-Peripheral Node | 1 | 0.00 | -1.11 |
| PPN | Ultra-Peripheral Node | 1 | 0.00 | -1.09 |
| VENT | Non-Hub Connector Node | 1 | 0.65 | -1.14 |
| CUN | Non-Hub Connector Node | 1 | 0.66 | -1.31 |
| B | Provincial Hub | 2 | 0.00 | 0.84 |
| VTA | Provincial Hub | 2 | 0.00 | 0.80 |
| DMH | Provincial Hub | 2 | 0.00 | 0.97 |
| PAG | Provincial Hub | 2 | 0.00 | 0.73 |
| VLPO | Provincial Hub | 2 | 0.00 | 0.75 |
| DP | Provincial Hub | 2 | 0.00 | 0.76 |
| LDT | Provincial Hub | 2 | 0.00 | 0.93 |
| PCG | Provincial Hub | 2 | 0.00 | 0.87 |
| SMT | Provincial Hub | 2 | 0.00 | 0.87 |
| PVHd | Provincial Hub | 2 | 0.00 | 1.00 |
| PT | Provincial Hub | 2 | 0.00 | 1.02 |
| PH | Provincial Hub | 2 | 0.00 | 1.09 |
| AHN | Provincial Hub | 2 | 0.00 | 0.97 |
| ILA | Provincial Hub | 2 | 0.00 | 0.88 |
| TRN | Provincial Hub | 2 | 0.02 | 1.11 |
| MBO | Provincial Hub | 2 | 0.03 | 0.97 |
| PVT | Provincial Hub | 2 | 0.03 | 0.88 |
| MPN | Provincial Hub | 2 | 0.03 | 0.97 |
| MD | Provincial Hub | 2 | 0.03 | 0.74 |
| LS | Provincial Hub | 2 | 0.05 | 1.00 |
| MPO | Provincial Hub | 2 | 0.05 | 0.88 |
| PALc | Provincial Hub | 2 | 0.15 | 0.74 |
| VMH | Provincial Hub | 2 | 0.17 | 0.74 |
| RE | Peripheral Node | 2 | 0.06 | 0.69 |
| TT | Peripheral Node | 2 | 0.06 | 0.28 |
| ACAd | Peripheral Node | 2 | 0.06 | 0.09 |
| AT | Peripheral Node | 2 | 0.11 | -0.12 |
| ACB | Peripheral Node | 2 | 0.15 | 0.34 |
| AM | Peripheral Node | 2 | 0.15 | 0.55 |
| PG | Peripheral Node | 2 | 0.16 | 0.06 |
| IV | Peripheral Node | 2 | 0.16 | -2.12 |
| AIv | Peripheral Node | 2 | 0.17 | -0.27 |

Supplementary Table 7 continued

| | | | | |
|---|---|---|---|---|
| PALd | Peripheral Node | 2 | 0.18 | 0.08 |
| PVH | Peripheral Node | 2 | 0.19 | 0.45 |
| CP | Peripheral Node | 2 | 0.19 | -2.24 |
| VTN | Peripheral Node | 2 | 0.21 | -1.86 |
| RCH | Peripheral Node | 2 | 0.22 | 0.29 |
| NI | Peripheral Node | 2 | 0.22 | 0.48 |
| LHA | Peripheral Node | 2 | 0.24 | 0.32 |
| SSp-ll | Peripheral Node | 2 | 0.24 | -1.97 |
| MSC | Peripheral Node | 2 | 0.24 | -0.35 |
| PRNc | Peripheral Node | 2 | 0.25 | -0.08 |
| III | Peripheral Node | 2 | 0.26 | -2.05 |
| RSP | Peripheral Node | 2 | 0.30 | -1.47 |
| AVP | Peripheral Node | 2 | 0.30 | -0.26 |
| PRT | Peripheral Node | 2 | 0.32 | -2.24 |
| ADP | Peripheral Node | 2 | 0.32 | -0.37 |
| DTN | Peripheral Node | 2 | 0.33 | -1.85 |
| LPO | Peripheral Node | 2 | 0.33 | -0.51 |
| MO | Peripheral Node | 2 | 0.35 | -1.39 |
| Ramb | Peripheral Node | 2 | 0.40 | -1.80 |
| SPA | Peripheral Node | 2 | 0.41 | -1.86 |
| CS | Peripheral Node | 2 | 0.41 | -1.97 |
| SSp-ul | Peripheral Node | 2 | 0.46 | -2.22 |
| PR | Ultra-Peripheral Node | 2 | 0.00 | 0.42 |
| ACAv | Ultra-Peripheral Node | 2 | 0.00 | 0.58 |
| PL | Ultra-Peripheral Node | 2 | 0.00 | 0.69 |
| MOB | Ultra-Peripheral Node | 2 | 0.00 | 0.16 |
| AON | Ultra-Peripheral Node | 2 | 0.00 | 0.35 |
| AOB | Ultra-Peripheral Node | 2 | 0.00 | -0.79 |
| SCs | Ultra-Peripheral Node | 2 | 0.00 | -0.45 |
| MRN | Ultra-Peripheral Node | 2 | 0.00 | -0.35 |
| PRNr | Ultra-Peripheral Node | 2 | 0.00 | -0.07 |
| SLC | Ultra-Peripheral Node | 2 | 0.00 | 0.23 |
| SLD | Ultra-Peripheral Node | 2 | 0.00 | 0.52 |
| ORBl | Ultra-Peripheral Node | 2 | 0.00 | -0.06 |
| PS | Ultra-Peripheral Node | 2 | 0.00 | 0.70 |
| RN | Ultra-Peripheral Node | 2 | 0.00 | -0.12 |
| ORBvl | Ultra-Peripheral Node | 2 | 0.00 | -0.11 |
| IAM | Ultra-Peripheral Node | 2 | 0.00 | 0.62 |
| EPI | Ultra-Peripheral Node | 2 | 0.00 | 0.67 |
| SCm | Ultra-Peripheral Node | 2 | 0.00 | -0.93 |
| ORBm | Ultra-Peripheral Node | 2 | 0.03 | 0.60 |
| PALv | Ultra-Peripheral Node | 2 | 0.03 | 0.28 |

Supplementary Table 7 continued

| | | | | |
|---|---|---|---|---|
| ILM | Ultra-Peripheral Node | 2 | 0.04 | -0.07 |
| EP | Connector Hub | 3 | 0.56 | 2.25 |
| MEA | Connector Hub | 3 | 0.59 | 1.74 |
| V | Connector Hub | 3 | 0.62 | 1.42 |
| VII | Connector Hub | 3 | 0.65 | 1.28 |
| AIp | Connector Hub | 3 | 0.66 | 0.83 |
| SNr | Peripheral Node | 3 | 0.24 | -0.37 |
| FN | Peripheral Node | 3 | 0.38 | -1.85 |
| PIR | Peripheral Node | 3 | 0.50 | -1.09 |
| VISC | Peripheral Node | 3 | 0.51 | -1.02 |
| BLA | Peripheral Node | 3 | 0.55 | 0.10 |
| LP | Peripheral Node | 3 | 0.55 | 0.61 |
| PSV | Peripheral Node | 3 | 0.56 | 0.40 |
| SO | Peripheral Node | 3 | 0.59 | 0.08 |
| IA | Peripheral Node | 3 | 0.59 | 0.35 |
| CA2 | Peripheral Node | 3 | 0.61 | 0.60 |
| AAA | Peripheral Node | 3 | 0.61 | 0.56 |
| LT | Peripheral Node | 3 | 0.61 | -1.58 |
| IRN | Non-Hub Connector Node | 3 | 0.63 | -0.53 |
| VP | Non-Hub Connector Node | 3 | 0.63 | -0.86 |
| BA | Non-Hub Connector Node | 3 | 0.63 | 0.61 |
| STN | Non-Hub Connector Node | 3 | 0.65 | -1.08 |
| AD | Non-Hub Connector Node | 3 | 0.65 | -0.56 |
| CA1 | Non-Hub Connector Node | 3 | 0.65 | -0.23 |
| SGN | Non-Hub Connector Node | 3 | 0.66 | -0.80 |
| LC | Non-Hub Connector Node | 3 | 0.66 | -0.44 |
| CENT | Non-Hub Connector Node | 3 | 0.66 | -0.44 |
| VISl | Provincial Hub | 4 | 0.00 | 0.93 |
| VISal | Provincial Hub | 4 | 0.00 | 1.36 |
| AUDv | Provincial Hub | 4 | 0.00 | 0.91 |
| AUDp | Provincial Hub | 4 | 0.00 | 1.24 |
| Tea | Provincial Hub | 4 | 0.00 | 1.28 |
| PRE | Provincial Hub | 4 | 0.26 | 0.72 |
| ECT | Peripheral Node | 4 | 0.61 | -0.53 |
| LGv | Ultra-Peripheral Node | 4 | 0.00 | -1.40 |
| POL | Ultra-Peripheral Node | 4 | 0.00 | -1.40 |
| SubG | Ultra-Peripheral Node | 4 | 0.00 | -1.40 |
| AUDd | Ultra-Peripheral Node | 4 | 0.00 | -1.02 |
| NB | Ultra-Peripheral Node | 4 | 0.00 | -1.00 |
| POST | Ultra-Peripheral Node | 4 | 0.00 | -0.45 |
| AUDpo | Ultra-Peripheral Node | 4 | 0.00 | -0.10 |
| VISp | Ultra-Peripheral Node | 4 | 0.00 | 0.41 |

Supplementary Table 7 continued

| | | | | |
|---|---|---|---|---|
| VISpl | Ultra-Peripheral Node | 4 | 0.00 | 0.43 |
| PO | Connector Hub | 5 | 0.38 | 0.76 |
| CUL | Connector Hub | 5 | 0.46 | 0.97 |
| ENT | Connector Hub | 5 | 0.46 | 1.25 |
| AN | Connector Hub | 5 | 0.48 | 1.16 |
| PFL | Connector Hub | 5 | 0.48 | 1.30 |
| CA3 | Connector Hub | 5 | 0.52 | 0.77 |
| PA | Connector Hub | 5 | 0.52 | 1.42 |
| LD | Connector Hub | 5 | 0.55 | 1.11 |
| TR | Connector Hub | 5 | 0.55 | 0.89 |
| VNC | Connector Hub | 5 | 0.57 | 0.87 |
| DG | Provincial Hub | 5 | 0.28 | 1.26 |
| IP | Peripheral Node | 5 | 0.17 | -0.33 |
| PBG | Peripheral Node | 5 | 0.36 | -1.30 |
| DN | Peripheral Node | 5 | 0.42 | -0.48 |
| SIM | Peripheral Node | 5 | 0.43 | 0.32 |
| SUB | Peripheral Node | 5 | 0.44 | 0.68 |
| PRM | Peripheral Node | 5 | 0.45 | -1.47 |
| IC | Peripheral Node | 5 | 0.47 | -1.46 |
| SOC | Peripheral Node | 5 | 0.48 | -0.30 |
| PARN | Peripheral Node | 5 | 0.49 | 0.47 |
| PST | Peripheral Node | 5 | 0.49 | -1.14 |
| PAA | Peripheral Node | 5 | 0.49 | -0.34 |
| COA | Peripheral Node | 5 | 0.50 | -0.66 |
| CN | Peripheral Node | 5 | 0.51 | -0.24 |
| PERI | Peripheral Node | 5 | 0.54 | 0.01 |
| PAR | Ultra-Peripheral Node | 5 | 0.00 | -1.77 |
| x | Ultra-Peripheral Node | 5 | 0.00 | -1.43 |
| NLL | Ultra-Peripheral Node | 5 | 0.00 | -1.31 |
| SPVO | Ultra-Peripheral Node | 5 | 0.00 | -0.96 |
| FL | Ultra-Peripheral Node | 5 | 0.00 | -0.07 |

*Supplementary Table 8.* Each brain region's module and classified role in the female frontloading functional connectivity network.

| Region | Category | Module | PC | WMDz |
|---|---|---|---|---|
| ORBl | Connector Hub | 1 | 0.48 | 1.24 |
| PERI | Peripheral Node | 1 | 0.19 | -0.99 |
| GU | Peripheral Node | 1 | 0.26 | 0.59 |
| AIv | Peripheral Node | 1 | 0.27 | 0.58 |
| FRP | Peripheral Node | 1 | 0.46 | 0.57 |
| AOB | Peripheral Node | 1 | 0.50 | -0.13 |
| PST | Ultra-Peripheral Node | 1 | 0.00 | -1.86 |
| PIR | Connector Hub | 2 | 0.33 | 0.98 |
| CEA | Connector Hub | 2 | 0.36 | 1.30 |
| FS | Connector Hub | 2 | 0.37 | 0.86 |
| EP | Connector Hub | 2 | 0.39 | 1.18 |
| PALv | Connector Hub | 2 | 0.42 | 1.33 |
| MEA | Connector Hub | 2 | 0.43 | 0.97 |
| BLA | Connector Hub | 2 | 0.44 | 1.19 |
| IA | Connector Hub | 2 | 0.45 | 1.11 |
| AAA | Provincial Hub | 2 | 0.30 | 0.99 |
| SSp-n | Peripheral Node | 2 | 0.12 | -0.79 |
| BA | Peripheral Node | 2 | 0.19 | -0.39 |
| AId | Peripheral Node | 2 | 0.19 | 0.49 |
| SSs | Peripheral Node | 2 | 0.21 | 0.22 |
| AIp | Peripheral Node | 2 | 0.21 | 0.44 |
| CLA | Peripheral Node | 2 | 0.26 | 0.23 |
| VISC | Peripheral Node | 2 | 0.27 | 0.10 |
| PAA | Peripheral Node | 2 | 0.34 | -1.07 |
| SSp-m | Peripheral Node | 2 | 0.35 | -0.39 |
| RR | Peripheral Node | 2 | 0.39 | -1.30 |
| CP | Peripheral Node | 2 | 0.40 | 0.70 |
| OT | Peripheral Node | 2 | 0.43 | -0.03 |
| PALd | Peripheral Node | 2 | 0.44 | 0.04 |
| STN | Peripheral Node | 2 | 0.45 | -1.17 |
| PO | Peripheral Node | 2 | 0.45 | -0.09 |
| SSp-tr | Peripheral Node | 2 | 0.45 | 0.15 |
| POL | Peripheral Node | 2 | 0.46 | -2.07 |
| SSp-ul | Peripheral Node | 2 | 0.48 | 0.11 |
| LD | Peripheral Node | 2 | 0.48 | 0.60 |
| MO | Peripheral Node | 2 | 0.48 | 0.58 |
| CA2 | Peripheral Node | 2 | 0.48 | -0.44 |
| SSp-ll | Peripheral Node | 2 | 0.49 | -0.04 |
| SAG | Peripheral Node | 2 | 0.49 | -1.84 |
| COA | Peripheral Node | 2 | 0.49 | 0.31 |
| SubG | Peripheral Node | 2 | 0.63 | -2.17 |

Supplementary Table 8 continued

| | | | | |
|---|---|---|---|---|
| ECT | Peripheral Node | 2 | 0.63 | -2.06 |
| CENT | Connector Hub | 3 | 0.54 | 0.98 |
| CN | Provincial Hub | 3 | 0.00 | 1.78 |
| CUL | Provincial Hub | 3 | 0.00 | 1.05 |
| IC | Provincial Hub | 3 | 0.00 | 1.75 |
| FL | Provincial Hub | 3 | 0.00 | 1.78 |
| SIM | Provincial Hub | 3 | 0.20 | 1.55 |
| PB | Peripheral Node | 3 | 0.27 | -0.76 |
| PSV | Peripheral Node | 3 | 0.38 | -0.13 |
| VISp | Peripheral Node | 3 | 0.41 | -0.10 |
| LC | Peripheral Node | 3 | 0.41 | -0.36 |
| PBG | Peripheral Node | 3 | 0.48 | -0.09 |
| VISal | Peripheral Node | 3 | 0.50 | -0.76 |
| POST | Peripheral Node | 3 | 0.50 | -1.08 |
| GENd | Peripheral Node | 3 | 0.50 | -0.75 |
| VISpl | Peripheral Node | 3 | 0.50 | -0.75 |
| SUT | Peripheral Node | 3 | 0.50 | -0.76 |
| VNC | Peripheral Node | 3 | 0.61 | -0.36 |
| SSp-bfd | Peripheral Node | 3 | 0.63 | -0.73 |
| PFL | Peripheral Node | 3 | 0.65 | 0.65 |
| SUB | Ultra-Peripheral Node | 3 | 0.00 | -1.08 |
| PPN | Ultra-Peripheral Node | 3 | 0.00 | -1.08 |
| LGv | Ultra-Peripheral Node | 3 | 0.00 | -0.73 |
| SNc | Connector Hub | 4 | 0.50 | 1.12 |
| CA3 | Connector Hub | 4 | 0.53 | 2.08 |
| LHA | Connector Hub | 4 | 0.54 | 0.80 |
| CA1 | Connector Hub | 4 | 0.56 | 1.46 |
| PA | Connector Hub | 4 | 0.57 | 0.95 |
| ILM | Connector Hub | 4 | 0.58 | 1.67 |
| DG | Connector Hub | 4 | 0.59 | 1.31 |
| Tea | Peripheral Node | 4 | 0.26 | -0.65 |
| AUDd | Peripheral Node | 4 | 0.31 | -0.88 |
| PP | Peripheral Node | 4 | 0.33 | -0.12 |
| SGN | Peripheral Node | 4 | 0.48 | -1.14 |
| VISam | Peripheral Node | 4 | 0.48 | -0.96 |
| VP | Peripheral Node | 4 | 0.50 | 0.68 |
| ZI | Peripheral Node | 4 | 0.53 | 0.63 |
| LP | Peripheral Node | 4 | 0.55 | -0.11 |
| SPF | Peripheral Node | 4 | 0.58 | 0.66 |
| SNr | Peripheral Node | 4 | 0.61 | -1.17 |
| VENT | Peripheral Node | 4 | 0.61 | -0.38 |
| ENT | Peripheral Node | 4 | 0.62 | -0.66 |
| TR | Peripheral Node | 4 | 0.64 | -0.64 |
| MRN | Peripheral Node | 4 | 0.64 | 0.60 |

Supplementary Table 8 continued

| | | | | |
|---|---|---|---|---|
| PAR | Peripheral Node | 4 | 0.65 | -0.67 |
| RSP | Peripheral Node | 4 | 0.74 | -0.37 |
| AUDpo | Ultra-Peripheral Node | 4 | 0.00 | -0.86 |
| NB | Ultra-Peripheral Node | 4 | 0.00 | -1.70 |
| AUDv | Ultra-Peripheral Node | 4 | 0.00 | -1.12 |
| AUDp | Ultra-Peripheral Node | 4 | 0.00 | -0.53 |
| RPO | Connector Hub | 5 | 0.35 | 1.22 |
| EW | Provincial Hub | 5 | 0.00 | 1.22 |
| VISl | Ultra-Peripheral Node | 5 | 0.00 | -0.82 |
| NTB | Ultra-Peripheral Node | 5 | 0.00 | -0.82 |
| MARN | Ultra-Peripheral Node | 5 | 0.00 | -0.82 |
| DMH | Provincial Hub | 6 | 0.00 | 1.06 |
| IAM | Provincial Hub | 6 | 0.00 | 0.90 |
| LS | Provincial Hub | 6 | 0.00 | 0.91 |
| AVP | Provincial Hub | 6 | 0.00 | 1.30 |
| AM | Provincial Hub | 6 | 0.00 | 0.72 |
| ILA | Provincial Hub | 6 | 0.00 | 0.81 |
| SMT | Provincial Hub | 6 | 0.00 | 0.84 |
| ORBm | Provincial Hub | 6 | 0.00 | 0.87 |
| PT | Provincial Hub | 6 | 0.00 | 0.93 |
| PVHd | Provincial Hub | 6 | 0.00 | 1.03 |
| RE | Provincial Hub | 6 | 0.00 | 1.17 |
| PVT | Provincial Hub | 6 | 0.00 | 0.81 |
| VMH | Provincial Hub | 6 | 0.00 | 0.81 |
| SF | Provincial Hub | 6 | 0.00 | 0.92 |
| PH | Provincial Hub | 6 | 0.04 | 1.20 |
| MPN | Provincial Hub | 6 | 0.04 | 1.25 |
| AHN | Provincial Hub | 6 | 0.05 | 0.70 |
| MSC | Provincial Hub | 6 | 0.08 | 1.06 |
| PS | Provincial Hub | 6 | 0.09 | 0.84 |
| MD | Provincial Hub | 6 | 0.15 | 0.98 |
| MBO | Provincial Hub | 6 | 0.16 | 0.72 |
| MPO | Peripheral Node | 6 | 0.05 | 0.50 |
| PVp | Peripheral Node | 6 | 0.06 | -0.25 |
| VLPO | Peripheral Node | 6 | 0.07 | -0.41 |
| PALc | Peripheral Node | 6 | 0.08 | -0.59 |
| AD | Peripheral Node | 6 | 0.09 | 0.42 |
| PL | Peripheral Node | 6 | 0.17 | -0.04 |
| TU | Peripheral Node | 6 | 0.17 | -1.55 |
| Ramb | Peripheral Node | 6 | 0.21 | 0.49 |
| ACAd | Peripheral Node | 6 | 0.23 | -0.30 |
| PAG | Peripheral Node | 6 | 0.27 | 0.56 |
| ACAv | Peripheral Node | 6 | 0.29 | 0.33 |
| EPI | Peripheral Node | 6 | 0.29 | 0.27 |

Supplementary Table 8 continued

| | | | | |
|---|---|---|---|---|
| ORBvl | Peripheral Node | 6 | 0.31 | -1.65 |
| VTA | Peripheral Node | 6 | 0.32 | 0.17 |
| RN | Peripheral Node | 6 | 0.39 | -1.68 |
| TRN | Peripheral Node | 6 | 0.42 | -1.37 |
| PG | Peripheral Node | 6 | 0.46 | -1.59 |
| AV | Peripheral Node | 6 | 0.55 | -1.27 |
| SFO | Ultra-Peripheral Node | 6 | 0.00 | -1.56 |
| IMD | Ultra-Peripheral Node | 6 | 0.00 | -1.47 |
| PVpo | Ultra-Peripheral Node | 6 | 0.00 | -1.15 |
| PD | Ultra-Peripheral Node | 6 | 0.00 | -1.82 |
| OV | Ultra-Peripheral Node | 6 | 0.00 | -1.57 |
| SH | Ultra-Peripheral Node | 6 | 0.00 | -1.34 |
| SCH | Ultra-Peripheral Node | 6 | 0.00 | -1.15 |
| AVPV | Ultra-Peripheral Node | 6 | 0.00 | -1.14 |
| SBPV | Ultra-Peripheral Node | 6 | 0.00 | -0.75 |
| PR | Ultra-Peripheral Node | 6 | 0.00 | 0.61 |
| TT | Ultra-Peripheral Node | 6 | 0.00 | 0.54 |
| RCH | Ultra-Peripheral Node | 6 | 0.00 | 0.55 |
| SPA | Ultra-Peripheral Node | 6 | 0.00 | -0.62 |
| MEPO | Ultra-Peripheral Node | 6 | 0.00 | -1.14 |
| ADP | Ultra-Peripheral Node | 6 | 0.00 | -0.95 |
| PVH | Ultra-Peripheral Node | 6 | 0.00 | 0.51 |
| AON | Connector Hub | 7 | 0.77 | 1.48 |
| ACB | Peripheral Node | 7 | 0.50 | -0.05 |
| MOB | Peripheral Node | 7 | 0.67 | -1.34 |
| LPO | Peripheral Node | 7 | 0.70 | -0.08 |
| CS | Connector Hub | 8 | 0.33 | 0.72 |
| NI | Connector Hub | 8 | 0.35 | 1.60 |
| LDT | Connector Hub | 8 | 0.41 | 1.67 |
| III | Provincial Hub | 8 | 0.16 | 1.17 |
| DTN | Provincial Hub | 8 | 0.25 | 1.63 |
| AT | Provincial Hub | 8 | 0.26 | 1.58 |
| VTN | Peripheral Node | 8 | 0.20 | 0.40 |
| VII | Peripheral Node | 8 | 0.38 | 0.27 |
| MEV | Peripheral Node | 8 | 0.43 | 0.04 |
| SO | Peripheral Node | 8 | 0.45 | -1.82 |
| SLC | Peripheral Node | 8 | 0.47 | -0.87 |
| VI | Peripheral Node | 8 | 0.47 | -0.88 |
| PCG | Peripheral Node | 8 | 0.48 | 0.65 |
| PRNc | Peripheral Node | 8 | 0.48 | 0.00 |
| PSTN | Peripheral Node | 8 | 0.50 | -0.91 |
| GRN | Peripheral Node | 8 | 0.52 | -0.18 |
| SLD | Peripheral Node | 8 | 0.53 | 0.36 |
| PARN | Peripheral Node | 8 | 0.55 | -0.79 |

Supplementary Table 8 continued

| | | | | |
|---|---|---|---|---|
| SCs | Peripheral Node | 8 | 0.56 | 0.06 |
| SCm | Peripheral Node | 8 | 0.63 | 0.37 |
| PRNr | Peripheral Node | 8 | 0.64 | -0.30 |
| B | Peripheral Node | 8 | 0.64 | 0.06 |
| SG | Peripheral Node | 8 | 0.64 | -0.81 |
| IRN | Peripheral Node | 8 | 0.67 | -0.43 |
| PRT | Peripheral Node | 8 | 0.69 | -0.62 |
| PRE | Ultra-Peripheral Node | 8 | 0.00 | -1.82 |
| V | Ultra-Peripheral Node | 8 | 0.00 | -1.82 |
| IV | Ultra-Peripheral Node | 8 | 0.00 | 0.68 |
| IP | Connector Hub | 9 | 0.35 | 0.72 |
| x | Peripheral Node | 9 | 0.16 | 0.21 |
| SPVO | Peripheral Node | 9 | 0.16 | 0.22 |
| PRM | Peripheral Node | 9 | 0.16 | 0.23 |
| FN | Peripheral Node | 9 | 0.36 | 0.63 |
| DN | Peripheral Node | 9 | 0.37 | 0.26 |
| PGRN | Peripheral Node | 9 | 0.42 | 0.45 |
| AN | Peripheral Node | 9 | 0.45 | -2.86 |
| PRP | Peripheral Node | 9 | 0.50 | 0.54 |
| ACVII | Peripheral Node | 9 | 0.53 | -0.41 |
